# Glial state changes and neuroinflammatory RIPK1 signaling are a key feature of ALS pathogenesis

**DOI:** 10.1101/2024.04.12.589201

**Authors:** Matija Zelic, Anna Blazier, Fabrizio Pontarelli, Michael LaMorte, Jeremy Huang, Ozge E. Tasdemir-Yilmaz, Yi Ren, Sean K. Ryan, Pavithra Krishnaswami, Mikhail Levit, Disha Sood, Yao Chen, Joseph Gans, Xinyan Tang, Jennifer Hsiao-Nakamoto, Fen Huang, Bailin Zhang, Giorgio Gaglia, Dimitry Ofengeim, Timothy R. Hammond

## Abstract

Amyotrophic lateral sclerosis (ALS) is a progressive neurodegenerative disease that causes motor neuron loss in the brain and spinal cord. Neuroinflammation driven by activated microglia and astrocytes is prominent in ALS, but an understanding of cell state dynamics and which pathways contribute to the disease remains unclear. Single nucleus RNA sequencing of ALS spinal cords demonstrated striking changes in glial cell states, including increased expression of inflammatory and glial activation markers. Many of these signals converged on RIPK1 and the necroptotic cell death pathway. Activation of the necroptosis pathway in ALS spinal cords was confirmed in a large bulk RNA sequencing dataset and at the protein level. Blocking RIPK1 kinase activity delayed symptom onset and motor impairment and modulated glial responses in SOD1^G93A^ mice. We used a human iPSC-derived motor neuron, astrocyte, and microglia tri-culture system to identify potential biomarkers secreted upon RIPK1 activation, inhibited pharmacologically *in vitro*, and modulated in the CSF of people with ALS treated with a RIPK1 inhibitor. These data reveal ALS-enriched glial populations associated with inflammation and suggest a deleterious role for neuroinflammatory signaling in ALS pathogenesis.

## Introduction

Amyotrophic lateral sclerosis (ALS) is a devastating, fatal neurodegenerative disease leading to progressive motor neuron loss and muscle weakness. There remains a large unmet medical need with only a few approved drugs with limited efficacy and an average patient survival of 3-5 years from symptom onset [1]. About 90% of ALS cases are sporadic, without any clear genetic cause. Genetic mutations identified in familial cases have suggested several pathologic mechanisms including mitochondrial damage, endoplasmic reticulum and oxidative stress, dysfunctional RNA metabolism, and protein aggregation, but causes of disease progression, especially in sporadic cases, remain unclear [1, 2].

The detrimental role of neuroinflammation in various neurodegenerative diseases, including ALS, is becoming more recognized. Microglial activation, emergence of reactive astrocytes, and immune dysfunction have been demonstrated in ALS mouse models and people with ALS [3–6]. Deletion of the mutant superoxide dismutase 1 (SOD1) transgene in either microglia or astrocytes extends survival by slowing disease progression but not onset in the SOD1^G93A^ mouse model of ALS [7–9], suggesting non-cell autonomous contributions to disease pathogenesis. For microglia, inhibition of CSF1R in SOD1^G93A^ mice delays disease progression and reduces motor neuron death [10]. Neurotoxic astrocytes are also present in ALS, and blocking the emergence of neurotoxic astrocytes by deleting the triggering immune factors in SOD1^G93A^ mice extends survival [6, 11]. Additionally, human astrocytes derived from people with sporadic or familial ALS can induce motor neuron death *in vitro* [12, 13].

Analysis of bulk RNA sequencing (RNA-seq) data from ALS motor cortex and spinal cord samples demonstrated significant increases in microglia and astrocyte marker gene expression as well as elevated expression of inflammation-related genes [14, 15]. However, whether specific subpopulations or states of glial cells emerge in disease and how to target those cells and accompanying neuroinflammation in ALS is still unclear. To address these questions, we performed bulk and single nucleus RNA sequencing (snRNA-seq) on cervical spinal cords from people with ALS and age- and sex-matched controls. There was an emergence of disease-associated microglia and astrocyte cell states, with prominent activation of inflammation-related genes in ALS spinal cords. The inflammation and cell death regulator receptor interacting protein kinase 1 (RIPK1) and the necroptotic cell death pathway, including downstream mediators receptor interacting protein kinase 3 (RIPK3) and the pseudokinase mixed lineage kinase domain-like (MLKL) [16, 17], were one of the main inflammation-related pathways dysregulated in ALS. We confirmed RIPK1 kinase upregulation in several sequencing datasets and at the protein level in human sporadic ALS spinal cord samples. To further explore and validate the role of neuroinflammation and RIPK1 in ALS, we showed that blocking the kinase activity of RIPK1 delayed symptom onset and motor impairment in SOD1^G93A^ mice and reduced neuroinflammatory responses. Using a human induced pluripotent stem cell (iPSC)-derived tri-culture system composed ofmotor neurons, microglia, and astrocytes [18], we generated a RIPK1-dependent gene and secretome signature, identifying several cytokines that are reduced following RIPK1 inhibitor administration. These cytokines were also elevated in the CSF of people with ALS and may be modulated with a CNS-penetrant RIPK1 kinase inhibitor. This patient sample and model-driven approach helped to unravel the complexity of glial activation and neuroinflammation in ALS and suggest that these pathways could be targeted in disease.

## Results

### Glial state changes and neuroinflammation are prominent in ALS spinal cords

ALS pathology is driven by complex interactions between neurons, glia, and immune cells that results in motor neuron death. There remains a poor understanding of cell-specific responses in the human ALS spinal cord, including differences in cell type composition, changes in cell state, and contribution of various cell types to neuroinflammation. To address these questions, we performed transcriptomic analysis of postmortem cervical spinal cord samples from 4 unaffected CNS controls and 8 people with ALS using bulk and snRNA-seq (Figure 1A). Bulk RNA-seq revealed broad gene expression changes in ALS spinal cords relative to controls, with gene ontology (GO) analysis demonstrating upregulation of inflammation and immune-related biological processes and downregulation of processes associated with synaptic signaling and neuron-related functions (Figures 1B-E). Various cytokines and inflammation-associated markers such as *SPP1*, *CHI3L2*, and *CCL2* were strongly upregulated in ALS spinal cords while decreased expression of *NEFL* and *CHAT* suggested neuronal loss (Figure 1D).

**Figure 1:**
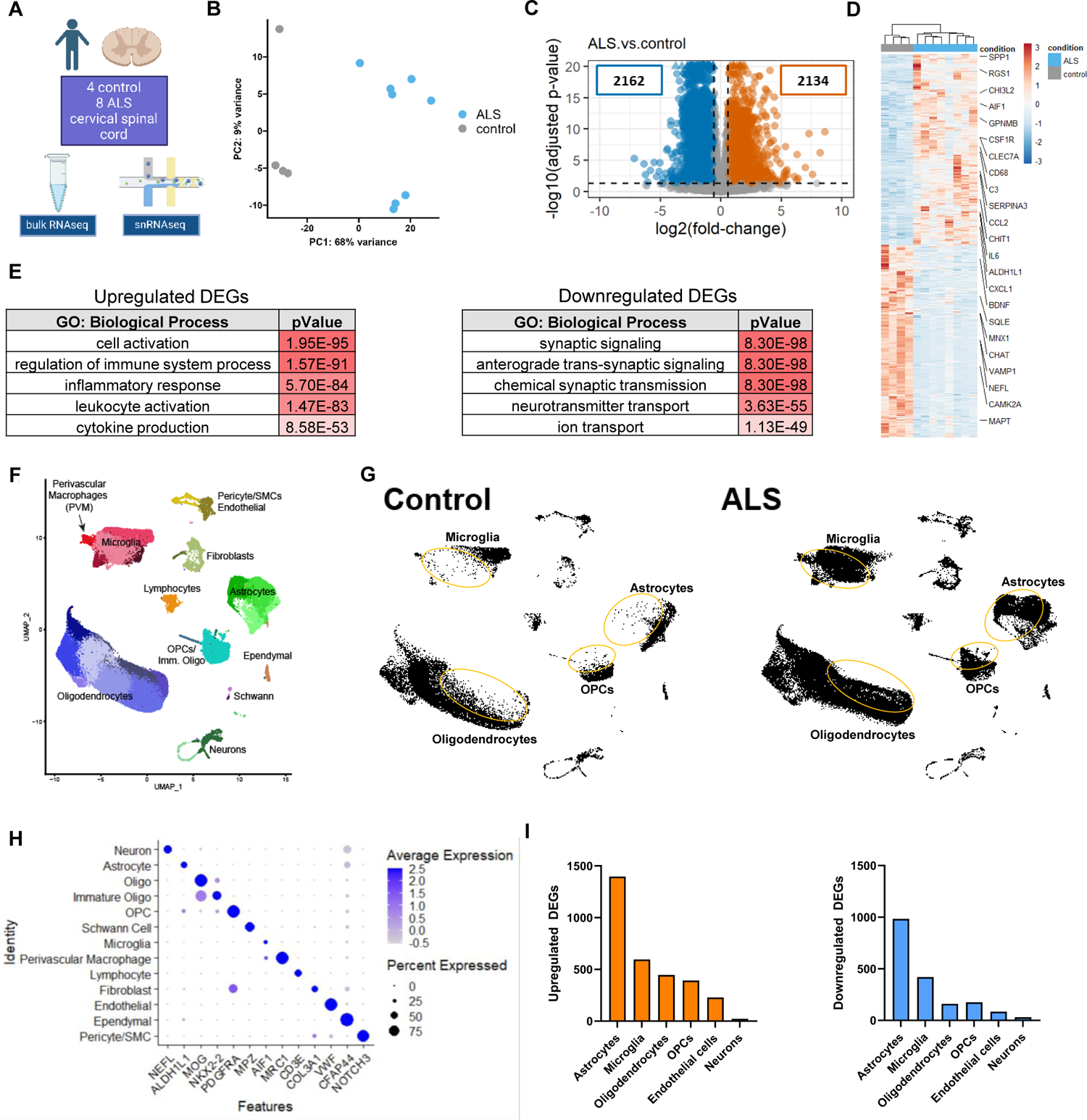
Inflammatory responses and glial state changes are prominent in ALS spinal cords A) Schematic of experimental approach for bulk and single cell RNA-seq of control (n=4) and ALS (n=8) postmortem spinal cord samples B) PCA of ALS and control spinal cords from bulk RNA –seq analysis C) Volcano plot showing DEGs in ALS versus control spinal cords (cutoff |FC| > 1.5, FDR < 0.05) D) Hierarchical heatmap clustering of DEGs in ALS spinal cord samples relative to controls E) Gene Ontology terms for biological processes from ToppGene Suite for upregulated and downregulated DEGs in ALS spinal cords relative to controls F) UMAP projection depicting all cells from snRNA-seq analysis of ALS and control spinal cords G) UMAP projection depicting all cells from either control or ALS spinal cords captured with snRNA-seq. Orange circles indicate subpopulations enriched in ALS spinal cord samples H) Dot plot showing the feature genes for all major cell types captured by snRNA-seq I) Bar graphs depicting total upregulated or downregulated DEGs from cell type-specific pseudobulk analysis of ALS and control spinal cords (cutoff |FC| > 1.5, FDR < 0.05) Data depict biological replicates. FDR (Benjamini-Hochberg) was used (E). See also Figures S1 and S2.

To assess transcriptional changes on the cellular level, we performed snRNA-seq on the same cervical spinal cord samples. Most of the major cell types were captured with the largest proportion from oligodendrocytes, microglia, and astrocytes (Figures 1F-H, and S1A-C). Consistent with the need for neuronal enrichment strategies, limited numbers of neurons were captured (Figures 1F, S1B and S1C) [19, 20]. Clustering of the cells revealed clear ALS-associated shifts in microglia, astrocytes, oligodendrocyte progenitor cells (OPCs), and oligodendrocytes (Figures 1G and S1D). Accounting for variance due to gender (Figures S2A and S2B), pseudobulk analysis of differentially expressed genes (DEGs) across all cells demonstrated enrichment of immune system and inflammation-related biological processes (Figures S2C-E), similar to the bulk RNA-seq analysis. Cell type-specific pseudobulk analysis revealed that the greatest number of DEGs were in astrocytes and microglia (Figure 1I). Together, this analysis demonstrates profound glial cell activation and cell state changes in ALS, suggesting dysregulation of neuroinflammatory signals may contribute to disease pathogenesis.

### ALS-enriched microglia and astrocytes adopt distinct transcriptional cell states

To further explore the nature of cell state dynamics in the top two cell types by DEGs, we performed subclustering analysis on microglia and astrocytes. Microglial subclustering revealed four distinct microglial populations (Mg 1 to Mg 4), with primarily Mg 2 and Mg 3 enriched in ALS spinal cords, but all subclusters expressed microglial genes such as *PTPRC* and *SPI1* (Figures 2A-C and S3A). The homeostatic Mg 1 cluster expressed high levels of microglial marker genes *CX3CR1* and *P2RY12* and was most represented in the control samples. These genes were strongly downregulated in the other microglial clusters (Figures 2B-D). The main ALS-enriched subpopulations had distinct enriched genes including immune (Mg 2: RNF144B, CXCR4) or neurodegenerative disease-associated markers (Mg 3: SPP1, HAMP) (Figure 2C). GO analysis demonstrated that relative to the homeostatic Mg 1 cluster, DEGs in the ALS-enriched microglial populations were involved in biological processes such as regulation of programmed cell death, inflammatory and immune system response, response to lipid, and response to hypoxia. Numerous genes were significantly upregulated in the microglia across the ALS patient samples, including those associated with inflammation like RGS1 and NAMPT [21, 22], lipid response genes such as ACSL1, which has recently been linked to lipid droplet laden microglia in disease [23], and disease-associated microglia (DAM) genes including APOE and FTH1 (Figures 2E-G) [24, 25]. While there was overlap of DEGs between the clusters (Figure S3B), potentially reflecting degrees of cell state activation, a balance of subset-specific and common microglial responses may exist in disease. Mg 2-specific DEGs were associated with regulation of lipid metabolic processes and response to hormone while Mg 3 DEGs were involved in oxidative phosphorylation and regulation of the immune system process. GO analysis of Mg 4 DEGs demonstrated biological processes related to protein folding and cellular responses to starvation or ER stress (Figures S3C and S3D). These gene expression changes show that microglia assume distinct cell states and upregulate various pathways in ALS that could contribute to neuroinflammation and play protective or damaging roles.

**Figure 2.**
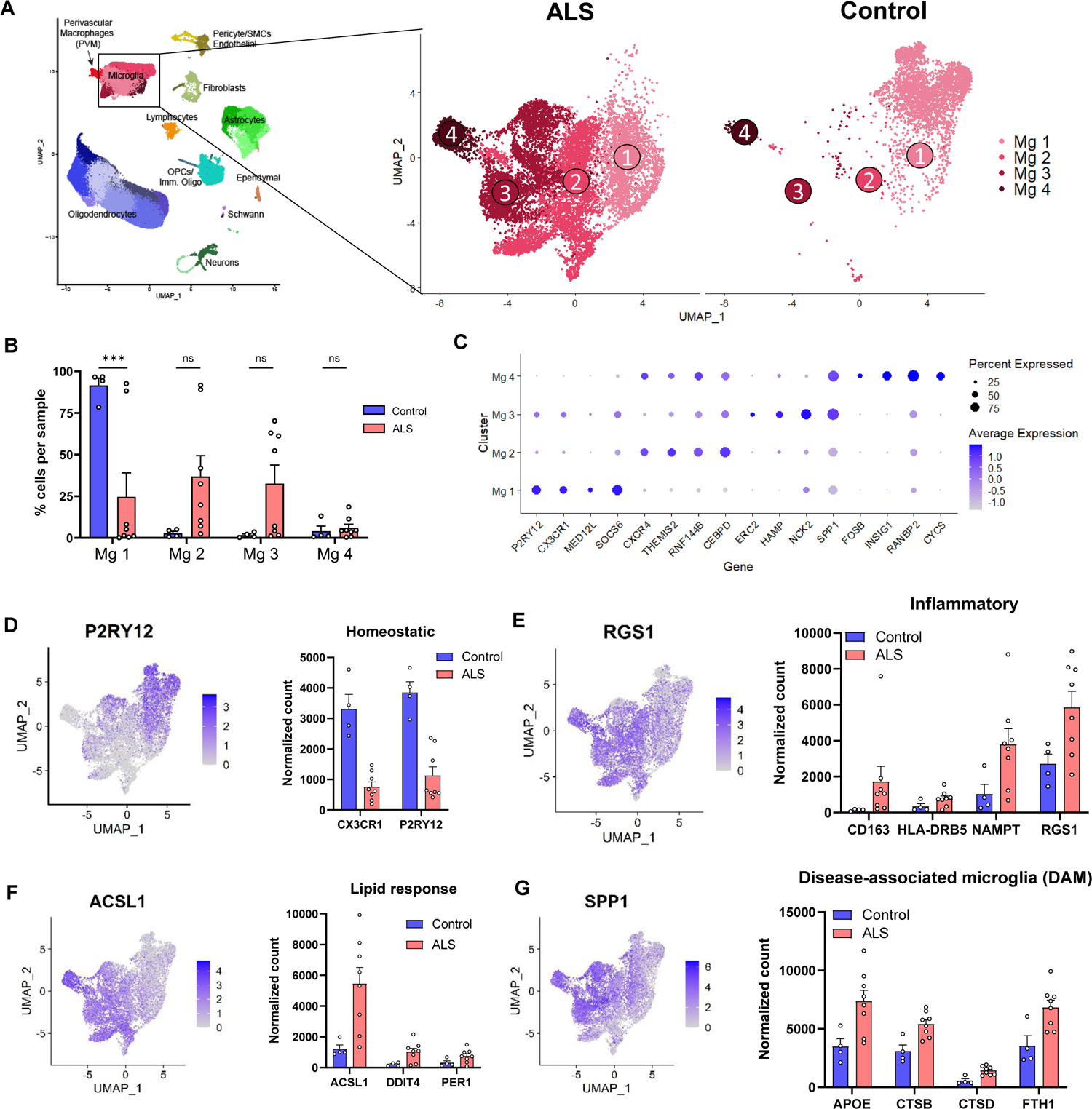
ALS spinal cord microglia adopt disease-associated cell states A) UMAP projection depicting subcluster analysis of ALS (n=8) and control (n=4) spinal cord microglia captured via snRNA-seq B) Quantification of microglia subcluster-specific contribution of cells per sample from ALS or control spinal cords C) Dot plot depicting the most significantly enriched genes per microglial subcluster D-G) UMAP projection of gene expression per cell and bar graph quantification of pseudobulk DESeq2 normalized counts highlighting example marker genes from homeostatic (D), inflammatory (E), lipid response (F), and DAM (G) biological processes in ALS (n=8) and control (n=4) spinal cord microglia (cutoff |FC| > 1.5, FDR < 0.05) Data depict biological replicates and error bars represent mean ± SEM. Two-way ANOVA with Sidak multiple comparisons test was performed (B). ns not significant, ^∗∗∗^p < 0.001. See also Figure S3.

In addition to the microglia, astrocytes were one of the most altered cell types in our dataset and have been shown to play a critical role in ALS [6, 9, 11]. Despite this, a clear signature of ALS-associated astrocytes has still not emerged. Astrocyte subclustering identified three distinct populations, with the Ast 1 cluster mainly present in the control samples while the Ast 2 and Ast 3 clusters were enriched in ALS spinal cords (Figures 3A and 3B). All astrocytes expressed canonical markers such as GFAP or SLC1A2, but unique DEGs defined each disease-associated cluster (Figures 3C, 3D, S4A and S4B). Expression of the transcription factor *BHLHE40*, a regulator of inflammation [26], was enriched in the Ast 2 cluster (Figures 3E and S4C). The chitinase genes *CHI3L1* and *CHI3L2*, reported as elevated in people with ALS and correlating with disease progression [27, 28], were significantly upregulated in the Ast 3 cluster (Figures 3C, 3E, and S4C). Pseudobulk analysis between control and ALS astrocytes demonstrated a broad upregulation of reactive astrocyte markers *EMP1*, *SERPINA3*, and *VIM*, as well as inflammatory cytokines and complement genes such as *CCL2* and *C3* (Figures 3D-F). These inflammation-related and reactive astrocyte genes were enriched in the Ast 3 cluster, with upregulated Ast 3 DEGs most significantly associated with ALS in the DisGeNET database of gene-disease associations (Figures 3D-F, S4C and S4D). These cell state shifts demonstrate that astrocytes undergo significant disease-associated changes and suggest a robust contribution by astrocytes to the neuroinflammatory environment in ALS.

**Figure 3.**
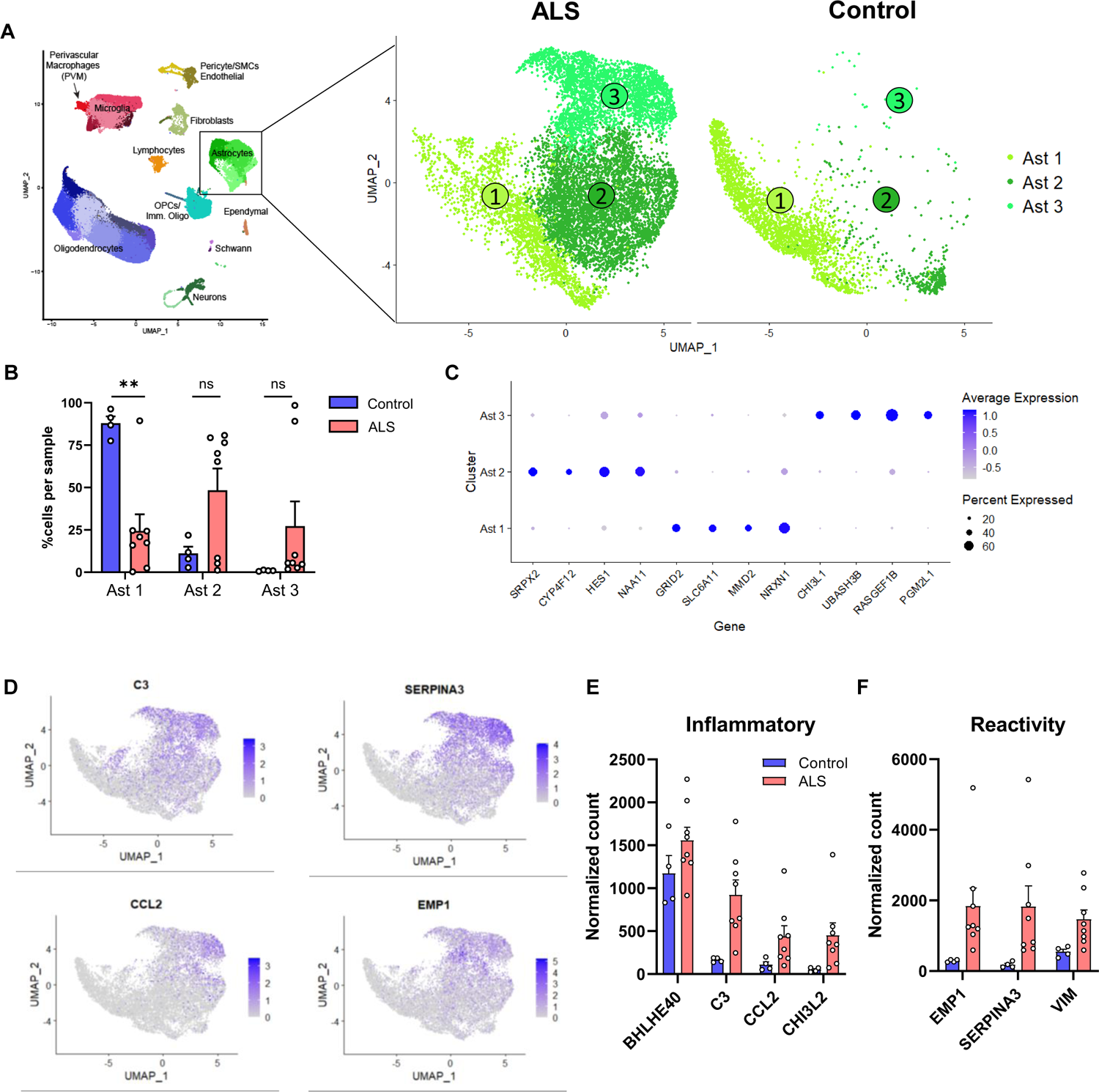
Astrocytes from ALS spinal cords transition to distinct subtypes with inflammatory and reactive transcriptomic responses A) UMAP projection depicting subcluster analysis of ALS (n=8) and control (n=4) spinal cord astrocytes captured via snRNA-seq B) Quantification of astrocyte subcluster-specific contribution of cells per sample from ALS or control spinal cords C) Dot plot depicting the most significantly enriched genes per astrocyte subcluster D-F) UMAP projection of gene expression per cell and bar graph of pseudobulk DESeq2 normalized counts highlighting inflammatory (D), and reactive (E) marker genes in ALS and control spinal cord astrocytes (cutoff |FC| > 1.5, FDR < 0.05) Data depict biological replicates and error bars represent mean ± SEM. Two-way ANOVA with Sidak multiple comparisons test was performed (B). ns not significant, ^∗∗^p < 0.01. See also Figure S4.

### RIPK1 expression and necroptosis signature are enriched in ALS spinal cords

Given the prominent inflammatory changes and alterations in microglial and astrocyte cell state accompanying motor neuron loss in ALS, we hypothesized that neuroinflammatory and/or cell death signaling hubs could be identified and potentially targeted in glial cells in disease. GO analysis of bulk RNA-seq DEGs identified the necroptotic signaling pathway, which includes RIPK1, as a significantly enriched cell death signaling node (Figure 4A). RIPK1, expressed in neurons and glia in the CNS, is a key regulator of both inflammation and cell death through its kinase activity [16, 29]. While several studies have associated RIPK1 kinase activation and necroptotic cell death with ALS pathology [30, 31], the role of RIPK1 signaling is not well understood [32, 33]. The expression of *RIPK1* and the downstream necroptosis mediators *RIPK3* and *MLKL* was significantly increased in our bulk RNA-seq dataset (Figure 4B). To expand the scope of necroptosis-related genes, we evaluated Qiagen’s Ingenuity Pathway Analysis (IPA) necroptosis gene list. Dysregulation of the necroptotic pathway in the bulk RNA-seq dataset was evident as the expression of 21 out of 34 necroptosis-related genes was significantly changed in ALS spinal cords relative to controls (Figure 4C). To further assess RIPK1 and necroptosis pathway activation in ALS, we analyzed the Target ALS bulk RNA-seq dataset across cervical, lumbar, and thoracic spinal cord samples from over 100 people with ALS. Relative to control spinal cords, ALS samples had substantial increases in *RIPK1*, *RIPK3*, and *MLKL* expression throughout the spinal cord (Figures 4D, 4E and S5A), establishing RIPK1 pathway upregulation as a feature of ALS pathology.

**Figure 4:**
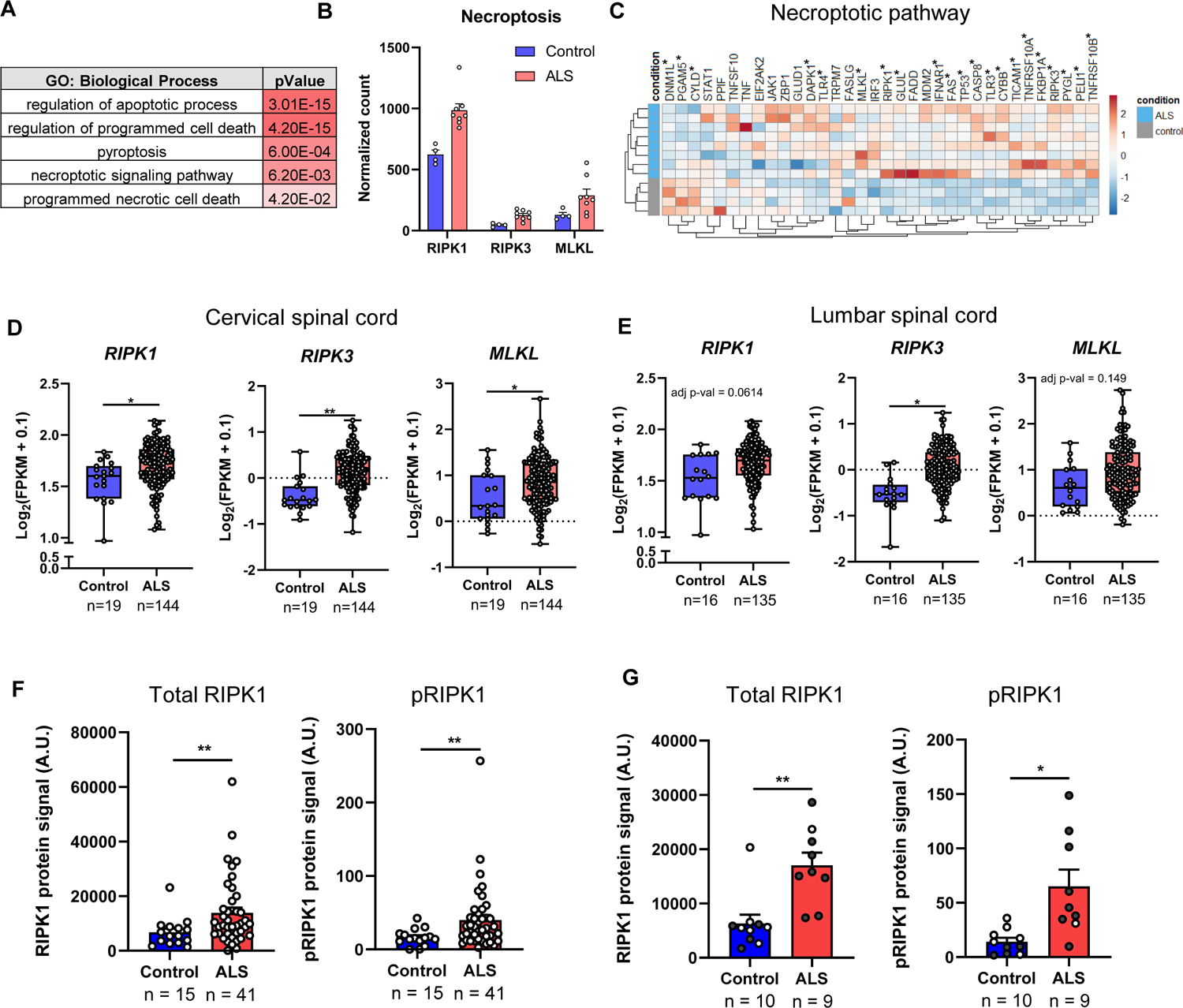
RIPK1 kinase activity and necroptosis pathway expression are increased in ALS postmortem spinal cord tissue A) Gene Ontology terms from ToppGene Suite for cell death-related biological processes for bulk RNA-seq DEGs in ALS spinal cords relative to control B) Gene expression levels of *RIPK1*, *RIPK3*, and *MLKL* in control (n=4) and ALS (n=8) spinal cords analyzed by bulk RNA-seq, depicting pseudobulk DESeq2 normalized counts (cutoff |FC| > 1.5, FDR < 0.05) C) Hierarchical heatmap clustering of the necroptosis pathway genes from IPA in ALS spinal cord samples relative to controls (FDR < 0.05) D, E) Box and whisker plots depicting gene expression levels of *RIPK1*, *RIPK3*, and *MLKL* in control and ALS cervical (D) and lumbar (E) spinal cords from the Target ALS bulk RNA-seq database F) MSD assay quantifying total and pRIPK1 levels in insoluble (RIPA-soluble) protein lysates from ALS (n=41) and control (n=15) spinal cords from Mass General Hospital and Target ALS cohorts G) MSD assay quantifying total and pRIPK1 levels in insoluble protein lysates from ALS (n=9) and control (n=10) cervical spinal cords samples. Black dots depict samples used for bulk and snRNA-seq Data depict biological replicates and error bars represent mean ± SEM. FDR (Benjamini Hochberg) in (A, D, and E) and unpaired two-tailed t test with Welch’s correction (F and G) were performed. ^∗^p < 0.05, ^∗∗^p < 0.01. See also Figures S5 and S6.

To determine whether RIPK1 is also increased at the protein level, we utilized the Meso Scale Discovery (MSD) platform to assess RIPK1 in spinal cords from a large cohort of people with primarily sporadic ALS. RIPK1 levels were elevated in ALS spinal cords, especially in the insoluble protein fraction, suggesting elevated RIPK1 kinase activity (Figures 4F and S5B, [29]). In the same tissue samples, we observed increased RIPK1 phosphorylation using a phospho-specific S166 RIPK1 antibody, confirming that the kinase activity of RIPK1 is significantly elevated in ALS spinal cords (Figures 4F and S5B). Increased RIPK1 activation and expression were observed in both cervical and thoracic spinal cord sections, as well as in the motor cortex (Figures 4F, 4G and S5B-E). We profiled total and pRIPK1 expression in our cohort of cervical spinal cord samples used for RNA-seq analysis. While there was a significant increase in both total and pRIPK1 levels in the ALS cervical spinal cord samples, the range in the overall expression (Figures 4G and S5E) suggested we may be able to correlate RIPK1 kinase activity to ALS-associated gene expression changes. Increased expression of microglial *CHI3L2*, the DAM marker *CTSD*, and the inflammatory cytokine *TNF* positively correlated with pRIPK1 levels while there was a negative correlation with the homeostatic microglial marker CX3CR1 (Figures S6A and S6B). In astrocytes, inflammation-related gene expression changes positively correlated with RIPK1 activation level, including the ALS-associated Ast 3-enriched *CCL2* and *CHI3L2* (Figures S6C and S6D), suggesting RIPK1 kinase activity may contribute to ALS-associated transcriptomic changes in glial cells. Overall, our data shows that RIPK1 expression and kinase activity are elevated in ALS spinal cords and may influence deleterious neuroinflammatory signaling in the disease.

### RIPK1 kinase inhibition delays disease progression and modulates neuroinflammation in SOD1^G93A^ mice

To better understand the role of neuroinflammation and RIPK1 signaling in ALS pathogenesis, we utilized the SOD1^G93A^ transgenic mouse model of ALS. Previous work suggested the involvement of RIPK1 and RIPK3 in this mouse model [30], but the overall importance of RIPK1 signaling and the necroptosis pathway to disease onset or progression remains unclear [30, 32–35]. Since there is variability in disease onset and progression in the SOD1 model with different genetic backgrounds, we utilized C57BL/6 mice, where symptom onset is delayed relative to the mixed B6/SJL background [36]. Starting on postnatal day 70, before noticeable motor deficits, mice were treated prophylactically with a CNS-penetrant RIPK1 kinase inhibitor [29]. The RIPK1 inhibitor was initially dosed in chow, resulting in good peripheral target engagement, prior to switching to administration via oral gavage when motor symptoms became more severe (Figures S7A-C). RIPK1 kinase inhibition delayed median symptom onset and weight loss in SOD1^G93A^ mice, resulting in a slight but insignificant extension in overall survival (Figures 5A-C and S7D). Plasma neurofilament levels were not affected by RIPK1 kinase inhibition (Figure S7E). To determine the effect of RIPK1 inhibition on motor impairment, we performed the wire hang and open field activity tests. While all WT and SOD1^G93A^ mice could initially hang suspended from the wire grid for the 3 minute cut-off, the performance of vehicle-treated SOD1^G93A^ mice declined rapidly. RIPK1 inhibition significantly delayed the latency to fall in the wire hang test (Figure 5D). The open field test was used to assess general locomotor activity for individual mice in a 20-minute period by measuring the overall distance traveled, speed of movement, and rearing counts as indicators of hind limb strength and exploratory behavior. Relative to WT mice, vehicle-treated SOD1^G93A^ mice declined in these parameters after 100 days of age while RIPK1 inhibition delayed motor impairment (Figures 5E-G). These data demonstrate a role for RIPK1 in contributing to disease progression and motor decline in the SOD1^G93A^ mouse model, supporting a deleterious role for RIPK1 and neuroinflammatory signaling in ALS pathogenesis.

**Figure 5:**
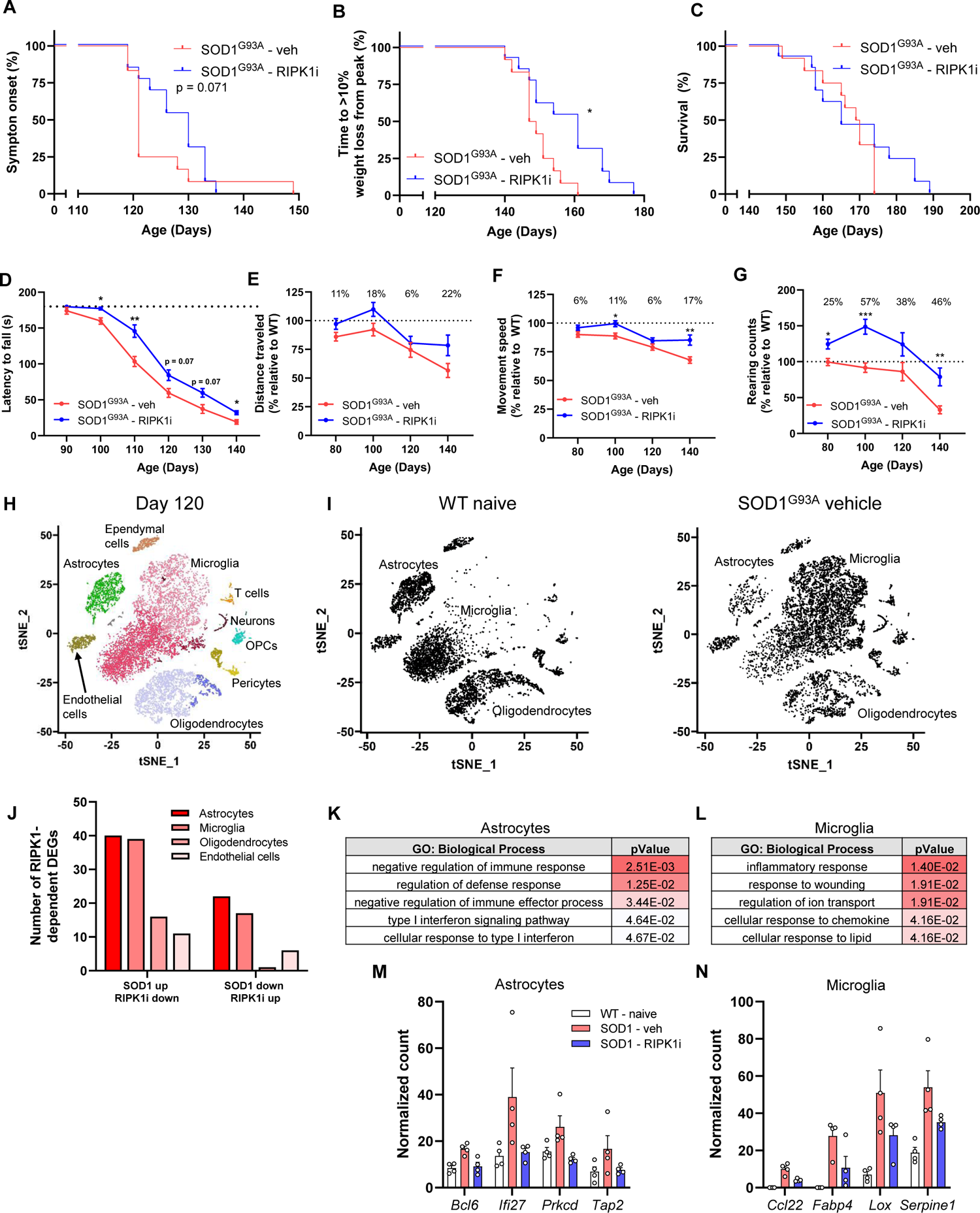
RIPK1 inhibition delays symptom onset and motor impairment in SOD1^G93A^ mice and modulates astrocyte and microglia gene expression A-C) Kaplan-Meier curves depicting symptom onset (A), weight loss from peak (B), and survival (C) in SOD1^G93A^ female mice treated with vehicle (n=12) or a CNS-penetrant RIPK1 kinase inhibitor (n=13), starting on postnatal day 70. D-G) Functional motor tests for wire hang (D) and open field activity parameters measuring distance traveled (E), movement speed (F), and rearing counts (G) in vehicle or RIPK1 inhibitor treated SOD1^G93A^ mice combined from two independent cohorts (n=10-24 per treatment group and time-point) H) tSNE plot depicting the major cell types captured by scRNA-seq from naïve WT, vehicle- and RIPK1 inhibitor-treated SOD1^G93A^ mice isolated on day 120 (n=4 per group) I) tSNE plots depicting all cells from either naïve WT or vehicle-treated SOD1^G93A^ mouse spinal cords from (H) J) Bar graph depicting total number of RIPK1-dependent DEGs in different cell types that were upregulated or downregulated in vehicle-treated SOD1^G93A^ mice relative to WT (cutoff |FC| > 1.5, p < 0.05) K, L) Gene Ontology terms for biological processes from ToppGene Suite that were increased in spinal cord astrocytes (K) or microglia (L) from vehicle-treated SOD1^G93A^ mice and reverted towards naïve WT levels with RIPK1 kinase inhibitor treatment M, N) Pseudobulk DESeq2 normalized counts highlighting example marker gene expression in astrocytes (M) and microglia (N) from the biological processes depicted in (K and L) for naïve WT, vehicle- and RIPK1 inhibitor-treated SOD1^G93A^ mouse spinal cords (cutoff |FC| > 1.5, p < 0.05) Data depict biological replicates and error bars represent mean ± SEM. Gehan-Breslow-Wilcoxon (A), Log-rank (B), two-way ANOVA with Sidak multiple comparisons test (D-G), or FDR (Benjamini-Hochberg) in (K, L) were performed. ^∗^p < 0.05, ^∗∗^p < 0.01, ^∗∗∗^p < 0.001. See also Figure S7.

To understand how RIPK1 inhibition may modulate cell signaling in SOD1^G93A^ mice, we performed single cell RNA-seq (scRNA-seq) on whole mouse spinal cords on postnatal day 120. This timepoint was selected because it is around symptom onset (Figures 5A and S7F) and was the period where improvements in motor function were initially observed with RIPK1 inhibition (Figures 5D-G). To generate cell suspensions, transcription inhibitors were included in the protease digestion and subsequent processing steps [37]. Glial cells were the most abundant cell types sequenced, with prominent microglial enrichment in vehicle-treated SOD1^G93A^ spinal cords relative to WT controls (Figures 5H, 5I and S7G-I). RIPK1 inhibition modulated dozens of SOD1^G93A^ gene changes in multiple glial cell types, especially astrocytes and microglia (Figure 5J). RIPK1 kinase-dependent DEGs were primarily upregulated in SOD1^G93A^ vehicle-treated mice and reduced towards homeostatic WT levels in RIPK1 inhibitor-treated SOD1^G93A^ mice (Figure 5J). RIPK1-dependent gene changes in astrocytes and microglia were associated with biological processes related to inflammation and regulation of immune response, type 1 interferon, and response to lipids (Figures 5K and 5L). RIPK1 kinase inhibition reduced immune and inflammation-related genes, including *Ifi27*, *Tap2*, *Fabp4*, and *Lox*, towards baseline levels in astrocytes and microglia (Figures 5M and 5N). Astrocytes upregulated *Ifi27* during cuprizone-induced demyelination [38] and *Tap2*, involved in antigen processing, is enriched in a subset of inflammatory astrocytes in the LPS neuroinflammation model [39]. *Fabp4*, a lipid binding protein, is involved in macrophage and microglia-mediated inflammation and drives cognitive decline in obese mice [40, 41]. *Lox* expression is increased in Alzheimer’s Disease and its inhibition may reduce parenchymal Aβ accumulation [42]. Altogether, these data demonstrate that RIPK1 kinase inhibition modulates gene expression in microglia and astrocytes, blocking increases in neuroinflammatory pathways that may contribute to the motor deficits observed in SOD1^G93A^ mice, thus suggesting that inhibiting RIPK1 could be a promising approach for the treatment of ALS.

### Identifying RIPK1 kinase-dependent changes in human motor neuron-glial tri-cultures

Increasingly, biomarkers of neurodegeneration and neuroinflammation are being used to track disease progression and assess efficacy in clinical trials for neurodegenerative diseases [43, 44]. While many of these biomarkers are still being validated, they are essential for trial success. To derive a human RIPK1-dependent inflammatory biomarker panel, we leveraged an iPSC-derived tri-culture system composed of ALS-relevant cell populations: motor neurons, astrocytes, and microglia (Figure 6A; [18]). All three cell types are critical since our recent work exploring the consequences of RIPK1 activation in mouse astrocytes and microglia demonstrated that RIPK1 kinase activity modulates inflammatory signaling *in vitro* and *in vivo* in both glial cell types [29]. To induce RIPK1 kinase activation, we stimulated the cells with TNF, Smac mimetic, and the pan-caspase inhibitor zVAD (TSZ) in the presence or absence of a RIPK1 kinase inhibitor (Figure 6A). To elucidate which tri-culture cells respond to RIPK1 activation and undergo transcriptional changes, we performed scRNA-seq (Figures S8A and S8B). TSZ stimulation caused a significant shift in the cell state of microglia and astrocytes, but not neurons, as reflected in the number of TSZ-modulated DEGs (Figures 6B and 6C). Pseudobulk analysis of the tri-culture DEGs demonstrated upregulation of immune and inflammatory genes that were regulated in a RIPK1 kinase-dependent manner (Figure 6D). To explain potential differences in the responsiveness of different tri-culture cell types to RIPK1 activation, we analyzed *RIPK1*, *RIPK3*, and *MLKL* expression. While *RIPK1* was highly expressed in both iPSC-derived microglia and astrocytes, its expression was much lower in neurons (Figure S8C). As we had previously observed in murine cells [29], *RIPK3* and *MLKL* were more highly expressed in microglia than astrocytes, albeit at lower expression levels compared to *RIPK1*. Neuronal expression of these genes was low, potentially explaining the general lack of neuronal transcriptional response to TSZ stimulation (Figures S8C-F). Therefore, RIPK1 activation in astrocytes and microglia is likely to drive inflammatory transcriptional responses, while neurons may respond at later time-points to signaling changes mediated by the glial cells. Cell-specific pseudobulk analysis demonstrated that astrocytes and especially microglia were the main drivers of the neuroinflammatory changes induced by RIPK1 activation in the tri-culture system (Figure 6E). Numerous cytokines and chemokines including *CCL3*, *IL1B*, and *CXCL1* were upregulated in a RIPK1 kinase-dependent manner in both cell types. RIPK1-regulated DEGs were sorted into biological processes related to immune cell migration and chemotaxis, inflammatory response, and response to lipids (Figures 6F-K). In neurons, RIPK1 kinase inhibition modulated several genes linked to immune response processes upon TSZ stimulation (Figures S8D-F). The astrocyte and microglia-mediated neuroinflammatory response driven by RIPK1 kinase activation in the iPSC tri-culture may represent a critical and conserved mechanism of RIPK1 signaling. The main biological processes upregulated in human ALS spinal cords and regulated in a RIPK1-dependent manner in astrocytes and microglia in SOD1^G93A^ mice were also related to immune and inflammatory responses, suggesting the relevance of targeting RIPK1 in deleterious neuroinflammatory processes that contribute to ALS pathogenesis.

**Figure 6:**
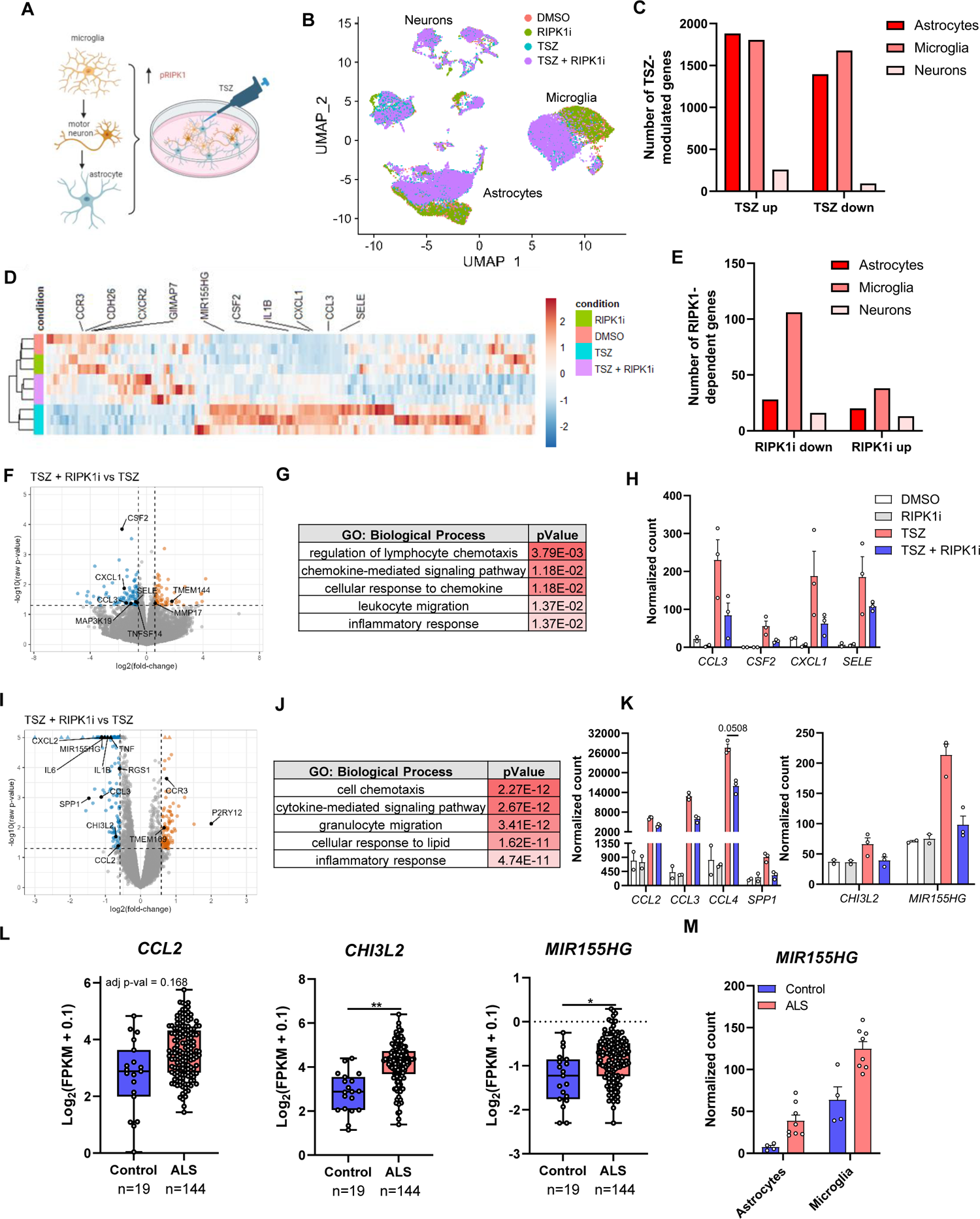
Astrocytes and microglia drive RIPK1-dependent neuroinflammatory changes in a human iPSC-derived tri-culture A) Schematic of experimental approach in iPSC tri-culture with TSZ stimulation, created with Biorender B) UMAP projection depicting all cells from scRNA-seq of tri-culture treated with TSZ in the presence or absence of a RIPK1 kinase inhibitor (RIPK1i) (n=2 DMSO, RIPK1i; n=3 TSZ, TSZ + RIPK1i) C) Bar graph depicting total upregulated or downregulated DEGs with TSZ stimulation from cell type-specific pseudobulk analysis of tri-culture (cutoff |FC| > 1.5, p < 0.05) D) Hierarchical heatmap clustering of DEGs modulated by RIPK1 kinase inhibition in TSZ-stimulated tri-culture cells assessed by pseudobulk analysis E) Bar graph depicting number of RIPK1-dependent DEGs reverted towards control levels from cell type-specific pseudobulk analysis of TSZ-stimulated tri-culture F, I) Volcano plot showing DEGs in astrocytes (F) or microglia (I) in the tri-culture samples stimulated with TSZ versus TSZ and RIPK1 inhibitor (TSZ + RIPK1i) G, J) Gene Ontology terms for biological processes for RIPK1-dependent genes upregulated with TSZ stimulation in tri-culture astrocytes (H) or microglia (J) H, K) Pseudobulk DESeq2 normalized count examples of inflammation-related RIPK1-dependent gene expression in TSZ-stimulated tri-culture astrocytes (H) or microglia (K) from the biological processes in (G) and (J) (cutoff |FC| > 1.5, p < 0.05) L) Gene expression levels of *CCL2*, *CHI3L2*, and *MIR155HG* in control and ALS cervical spinal cords from the Target ALS bulk RNA-seq database M) Pseudobulk DESeq2 normalized counts for *MIR155HG* gene expression in astrocytes and microglia from snRNA-seq of human ALS (n=8) and control (n=4) cervical spinal cords (from Figure 1) Data depict biological replicates (L) or technical replicates from a tri-culture, with two individual wells pooled for each technical replicate, and error bars represent mean ± SEM. FDR (Benjamini-Hochberg) was used (G, J, and L). ^∗^ p < 0.05, ^∗∗^p < 0.01; T, TNF; S, Smac mimetic; Z, zVAD-fmk. See also Figure S8.

Microglia from the stimulated tri-culture demonstrated RIPK1 kinase-dependent increases in *CCL2*, *CHI3L2*, and *MIR155HG* (Figure 6K). Chitinase genes have been linked to ALS progression and disease duration, and *MIR155HG* is a microRNA suggested to be overactive in the microglia of people with ALS, contributing to increased inflammation [27, 45]. Increased expression of these genes was observed in ALS cervical spinal cords in the Target ALS bulk RNA-seq dataset (Figure 6L), and more specifically in the microglia and astrocytes from our snRNA-seq analysis (Figures 3E and 6M). Identification of target- or disease-relevant biomarkers is important for patient stratification and understanding disease state and progression. Having profiled RIPK1 activation and expression levels in human spinal cord samples, we can assess RIPK1 correlation of ALS spinal cord DEGs with potential RIPK1-regulated biomarkers derived from the iPSC tri-culture. The transcriptional regulation of *CCL2*, *CHI3L2*, and *MIR155HG* in the tri-culture system is aligned with the astrocyte and microglia correlation of gene expression and amount of RIPK1 kinase activation in our ALS spinal cord snRNA-seq data (Figures S6B and S6D), validating the translational relevance of our RIPK1-dependent tri-culture gene signature.

To further explore these RIPK1-dependent neuroinflammatory genes as potential biomarkers, we performed Olink proteomic analysis of the tri-culture supernatants (Figure 7A). Several RIPK1-dependent cytokines from the single cell analysis, including CCL3, CCL4, and IL1B, were secreted upon TSZ stimulation and reversed by the RIPK1 inhibitor (Figure 7B). Studies have shown that these cytokines are elevated in the plasma and CSF of people with ALS relative to controls [3, 46–48]. We validated increased levels of multiple cytokines, including CCL2 and CCL3, in 9 ALS versus 9 control CSF samples (Figures 7C and 7D), demonstrating the translational relevance of these RIPK1-dependent biomarkers. In a Phase 1b clinical trial (NCT03757351) with a CNS-penetrant RIPK1 inhibitor [49], dosing with SAR443060 for 29 days generally reduced CCL2 and CCL4 levels in the CSF of people with ALS relative to the placebo dosing period (Figures 7E-I). These trends aligned with the level of RIPK1 target engagement in the CSF as assessed by an epitope masking assay (Figures 7G and 7I) [50]. Together, these data identify RIPK1-dependent biomarkers that are dysregulated in ALS and could be used in the clinic to monitor neuroinflammatory pathway activation and drug treatment effect. Much larger cohorts in additional clinical trials are needed to validate these biomarkers and assess the potential efficacy of RIPK1 inhibition in ALS. Beyond RIPK1, our data identify a neuroinflammatory node driven by unique cell state changes in microglia and astrocytes that must be studied further to understand how different glial subpopulations contribute to disease progression.

**Figure 7:**
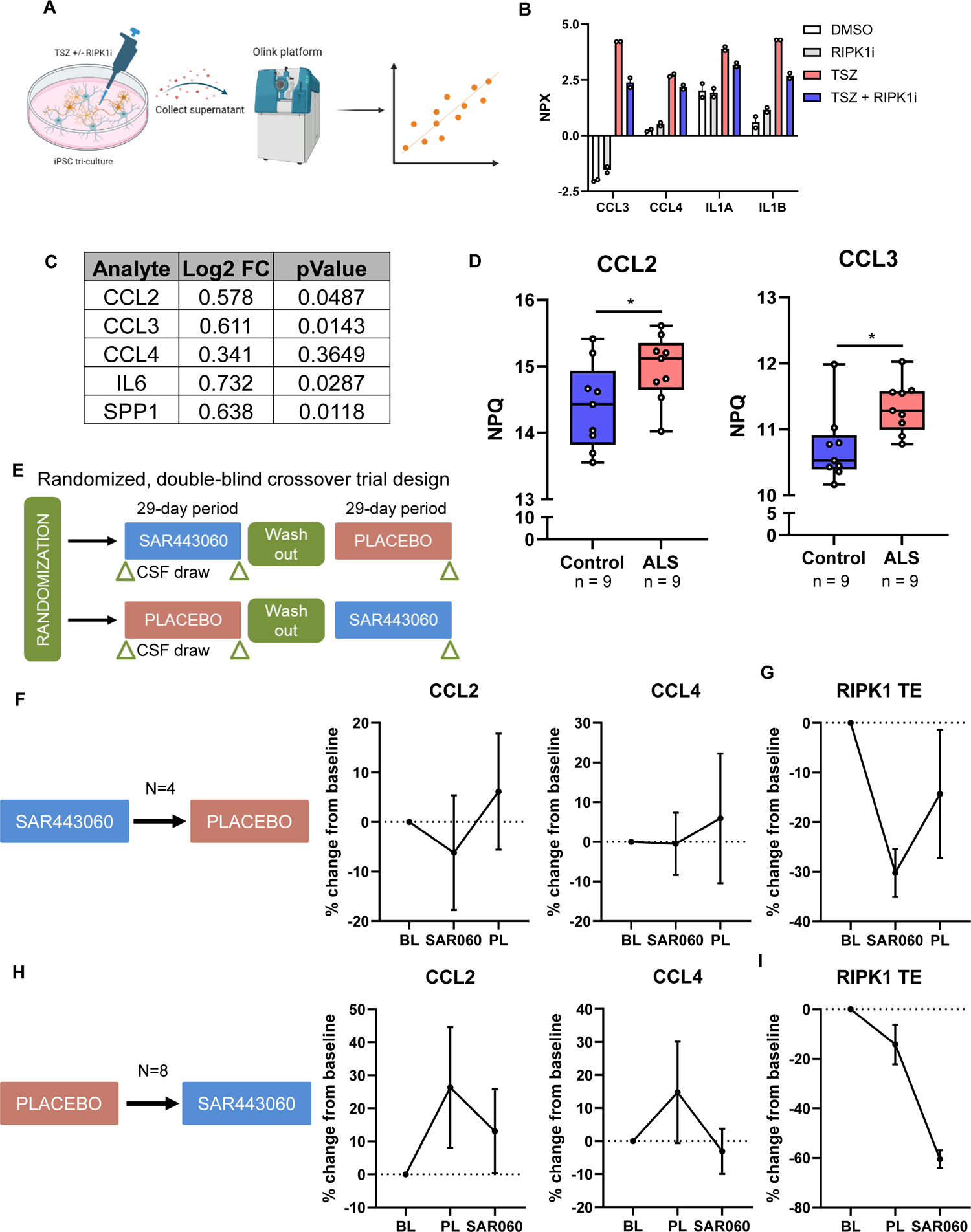
Identification of RIPK1-modulated biomarkers in ALS CSF samples A) Schematic of experimental approach for tri-culture supernatant analysis with Olink platform, created with Biorender B) Bar graph depicting Olink normalized protein expression of analytes from tri-culture stimulated with TSZ in the presence or absence of RIPK1 inhibitor (n=2) with cutoff |NPX| > 0.5, p < 0.05 C) Table depicting overlap of RIPK1-dependent tri-culture DEGs and analytes significantly enriched in ALS CSF samples measured on the NULISA platform D) Quantification of CCL2 and CCL3 levels in control and ALS CSF from (C) E) Design of Phase 1b trial (NCT03757351) with RIPK1 inhibitor (SAR443060) and placebo dosing in people with ALS F-I) Change in CCL2 and CCL4 levels (F, H) and RIPK1 target engagement (G, I) relative to baseline in the CSF of people with ALS dosed as depicted in (E) with n=4 (F) and n=8 (H) Data depict biological replicates or technical replicates from individuals wells in the tri-culture (B) and error bars represent mean ± SEM. One-tailed (B) or two-tailed (D) t tests were performed. ^∗^ p < 0.05; BL, baseline; PL, placebo; SAR060, SAR443060

## Discussion

We demonstrate that sporadic ALS-associated changes in human postmortem spinal cord samples are most prominent in astrocytes and microglia, including the emergence of disease-specific cell states and altered neuroinflammatory and immune-mediated signaling. Furthermore, necroptosis was identified as a significantly changed signaling node and we show that RIPK1 kinase activation and expression are elevated in ALS spinal cords, suggesting excessive RIPK1 signaling may contribute to the ALS-associated neuroinflammatory environment. RIPK1 inhibition delays symptom onset and motor impairment in SOD1^G93A^ mice and RIPK1 kinase activation in an iPSC-derived tri-culture system drives astrocyte and microglia-mediated inflammatory signaling, including changes in clinically relevant RIPK1-dependent biomarkers. Overall, our results demonstrate that aberrant RIPK1 kinase-dependent signaling may contribute to ALS pathogenesis by causing excessive neuroinflammation.

While the detrimental role of neuroinflammation in neurodegenerative diseases such as ALS is becoming more established [1, 3, 5, 6, 15, 51], how glial cell states are altered and contribute to disease is less clear. Two recent studies analyzed bulk RNA-seq data from over 150 ALS postmortem motor cortex and spinal cord samples and demonstrated significant increases in microglia and astrocyte marker gene expression, suggesting pro-inflammatory changes driven by these glial cells in ALS [14, 15]. However, the lack of single cell resolution and cell type understanding that can be obtained from bulk RNA-seq datasets limits the utility and strength of these analyses. In this study, we perform bulk and single nucleus RNA-sequencing on control and ALS cervical spinal cord samples, providing a key resource to the scientific community allowing for analysis of disease-associated changes on specific cell types of interest. Our analyses confirm and extend previous findings [14, 15, 51–53], demonstrating significant upregulation of an immune-related and pro-inflammatory gene signature in ALS spinal cords driven by disease-associated changes in microglia and astrocytes. Microglial subclustering revealed a loss of homeostatic microglial markers and enrichment of inflammation- and lipid-related genes, along with an increase in DAM markers such as *APOE* and *CTSD*, in ALS spinal cords. ALS-enriched astrocyte clusters upregulated activation markers such as *SERPINA3* and inflammatory markers including the chitinase-like proteins *CHI3L1* and *CHI3L2*, the complement component *C3*, and the cytokine *CCL2*. *CHI3L1* and *CHI3L2* are reported to be elevated in the motor cortex and spinal cord of people with sporadic ALS and correlated with shorter disease duration [27, 28]. Our transcriptomic data confirmed these findings and demonstrated that subsets of ALS-enriched astrocytes undergo this reactive cell state change. It is possible that there were differences in the level of motor neuron degeneration or neuroinflammation throughout the tissue samples. Thus, it is unclear if the different microglial and astrocyte states represent unique responses or a continuum of activation captured across areas with varying levels of pathology. However, there is an enrichment of inflammatory microglia and astrocyte cell states across the ALS spinal cords and many of these markers were conserved in larger patient datasets, including Target ALS, suggesting that these states may exist broadly in disease. Whether different subpopulations are physically associated with damaged neurons or other pathological features will be interesting to explore in a larger patient cohort.

RIPK1 emerged as a key neuroinflammatory signaling node in our ALS spinal cord sequencing data. RIPK1 kinase activity can drive both pro-inflammatory signaling and cell death, namely RIPK1 kinase-dependent apoptosis or caspase-independent necroptosis via RIPK3 and MLKL [16, 17]. Several genes implicated in familial ALS, including *TBK1* and *OPTN*, are negative regulators of RIPK1 signaling. Genetic and pharmacological inactivation of RIPK1 rescues elevated RIPK1 activity and necroptosis in the spinal cords of *Optn*^-/-^ mice [30]. Additionally, TBK1 suppresses RIPK1 activation by phosphorylation at T189 and the embryonic lethality of *Tbk1^-/-^* mice is rescued by genetic inactivation of RIPK1 kinase activity [31]. We had previously identified that RIPK1 drives a deleterious neuroinflammatory transcriptional signature in murine astrocytes and microglia and demonstrated that RIPK1 kinase-dependent signaling may contribute to MS pathology, particularly in progressive MS [29]. RIPK1 and the necroptotic cell death pathway have also been implicated in ALS mouse models and postmortem samples from people with ALS [30, 31, 34], but several studies did not replicate these findings [32, 33, 35], leaving the role of RIPK1 in ALS unclear. To functionally assess this potential target, we analyzed RIPK1 activation and expression at the mRNA and protein levels in several human tissue cohorts composed of primarily sporadic ALS samples. Consistent with the literature [32], we did not find a significant increase in RIPK1 expression in ALS motor cortex samples, suggesting that RIPK1 signaling may not be involved in upper motor neuron loss and motor cortex pathology. The potential difference of RIPK1 expression in upper and lower motor neurons, and perhaps differential susceptibility to necroptosis, needs to be investigated further. However, the expression of the core necroptosis components *RIPK1*, *RIPK3*, and *MLKL* and the broader necroptosis pathway genes were increased in ALS cervical spinal cords relative to controls in our bulk RNA-seq analysis. We confirmed a significant upregulation of *RIPK1*, *RIPK3*, and *MLKL* expression by analyzing Target ALS bulk RNA-seq data from over 100 ALS spinal cord samples across cervical, lumbar, and thoracic regions. Additionally, we demonstrated increased RIPK1 kinase activation and expression at the protein level throughout the spinal cord, including in cervical and thoracic ALS samples. Unlike previous studies which used traditional Western blotting [32, 35], we utilized the increased sensitivity of the MSD platform to assess RIPK1 protein levels. Furthermore, we used multiple tissue cohorts to analyze a much larger number of samples than previous studies [32, 33, 35]. While we were not able to correlate RIPK1 levels to disease duration or location of disease onset, this could be due to limitations in available patient meta-data or insufficient sample size across tissue regions or disease subtypes. However, the significant increase in RIPK1 RNA and protein expression in sporadic ALS spinal cords suggests this target may be relevant to both familial and sporadic forms of ALS [30, 31, 34, 54, 55].

Although we did not assess the role of RIPK3 or MLKL, we confirmed that pharmacologic inhibition of RIPK1 kinase activity with a CNS-penetrant RIPK1 inhibitor can delay symptom onset in the SOD1^G93A^ mouse model of ALS [30]. Additionally, we demonstrated that RIPK1 inhibition can delay motor impairment. While there was efficacy with RIPK1 inhibition in SOD1^G93A^ mice, motor impairment was modestly delayed and overall survival was not significantly affected. Higher sustained target inhibition may afford a more beneficial effect as mice dosed with the RIPK1 inhibitor in chow only ate ∼80% of the expected amount and target engagement decreased throughout the day. Alternatively, modulating inflammation in SOD1^G93A^ mice may only have a moderate effect or other inflammatory and cell death pathways, such as complement or ferroptosis, may contribute to disease progression [11, 56]. This may not be unexpected given the overexpression of the mutant SOD1 transgene. While there were hundreds of DEGs in the different glial cell populations in SOD1^G93A^ vehicle-treated mice compared to WT, less than 10% of the mouse DEGs were similarly changed in our human ALS spinal cord dataset (Figure S7J). Although the SOD1^G93A^ mouse is one of the most frequently used ALS mouse models, the broad transcriptomic changes observed with transgenic overexpression of this familial ALS risk gene may not be reflective of end-stage transcriptomic changes observed in human postmortem ALS spinal cords. While our scRNA-seq data could not confirm the existence of RIPK1-regulated inflammatory microglia (RRIMs) [34], we did observe RIPK1-regulated transcriptional effects on astrocytes and microglia related to immune and inflammatory responses. Overall, our data confirm a detrimental role for RIPK1 kinase activity in SOD1^G93A^ mice but suggest that RIPK1 signaling may be more prominent in mouse models with familial mutations associated with RIPK1 kinase regulation, such as *OPTN* and *TBK1* [30, 31].

Since our recent work explored the effect of RIPK1 activation in mouse astrocytes and microglia [29], we utilized an iPSC-derived tri-culture system containing motor neurons [18] to derive a human RIPK1-dependent gene signature and gain insight into potential biomarkers. RIPK1 kinase activation drove inflammation-related signaling changes in astrocytes and especially microglia, consistent with previous studies demonstrating these glial cells are the main mediators of inflammatory RIPK1-dependent signaling [29, 57, 58]. Interestingly, we identified microglial upregulation of *CCL2*, *CHI3L2*, and *MIR155HG* in our tri-culture system. These inflammatory regulators and potential biomarkers have been found elevated in SOD1^G93A^ mice and spinal cord tissue from people with familial and sporadic ALS, with *CHI3L2* correlated with shorter disease duration [3, 27, 45]. At the protein level we observed RIPK1-dependent regulation of cytokines such as CCL3, CCL4 and IL1B which are elevated in the plasma and CSF of people with ALS [3, 46–48, 59, 60]. In a Phase 1b clinical trial, both CCL2 and CCL4 were partially modulated in the CSF of people with ALS treated with the CNS-penetrant SAR443060 RIPK1 inhibitor relative to the placebo treatment period, although median inhibition of peripheral pRIPK1 at trough was only 66% [49]. Longer treatment duration or sustained peripheral and CNS RIPK1 kinase inhibition higher than that achieved by SAR443060 may allow for a greater impact on neuroinflammatory signaling. Overall, these data support the value of the human iPSC-derived tri-culture system to identify disease-relevant transcriptomic signatures and potential biomarkers that could be applicable for patient stratification in clinical trials.

Increased RIPK1 activation observed in ALS spinal cords may not cause pathogenic changes but rather be an indirect response to end-stage disease. However, the genetic association of negative RIPK1 regulators with familial ALS, elevated RIPK1 expression at both mRNA and protein levels in multiple tissue cohorts across spinal cord regions, efficacy of RIPK1 inhibition in SOD1^G93A^ mice, and translational relevance of the RIPK1-dependent transcriptional signature in the iPSC-derived tri-culture system suggest that aberrant RIPK1 kinase activation contributes to ALS pathogenesis. To what extent other inflammatory and pro-death signaling pathways influence disease or whether reducing neuroinflammation can be a disease-modifying treatment in ALS remain open questions. RIPK1 levels were recently reported to be increased in ALS serum, especially in people with bulbar onset ALS, suggesting potential utility as a peripheral biomarker and tool for patient stratification in clinical trials [61]. Our human ALS snRNA-seq data demonstrated a positive correlation of spinal cord RIPK1 activation with microglia and astrocyte-related genes such as *CCL2, CHI3L2*, and *TNF*, suggesting RIPK1 kinase activity contributes to inflammatory glial activation in ALS. Overall, our data demonstrate RIPK1 kinase-dependent signaling in both astrocytes and microglia can mediate neuroinflammation and may contribute to ALS pathogenesis, suggesting RIPK1 as a therapeutic target in ALS.

---

**Table S1:** Control and ALS cervical spinal cord samples used for RNA sequencing Samples in bold were used for sequencing from this sample cohort. RIN: RNA integrity number, PMI: postmortem interval.

| Donor ID | Diagnosis | Age | Sex | RIN | PMI (h) |
| --- | --- | --- | --- | --- | --- |
| **13085** | Unaffected control | 57 | Male | 8.9 | 22.35 |
| 13106 | Unaffected control | 59 | Male | 9.6 | 5.85 |
| 13107 | Unaffected control | 61 | Male | 9.5 | 16.65 |
| **13122** | Unaffected control | 68 | Female | 8.7 | 26.5 |
| 13170 | Unaffected control | 59 | Male | 8.9 | 11.65 |
| 13267 | Unaffected control | 58 | Female | 8.8 | 10.43 |
| **13394** | Unaffected control | 65 | Female | 9 | 16.82 |
| 13440 | Unaffected control | 68 | Male | 8.6 | 17.1 |
| **13446** | Unaffected control | 59 | Female | 7.1 | 17.47 |
| 13591 | Unaffected control | 71 | Male | 9.4 | 27.87 |
| **5724** | ALS | 65 | Male | 7.5 | 25 |
| **5781** | ALS | 61 | Female | 7.2 | 3 |
| **5867** | ALS | 51 | Male | 7.8 | 19 |
| **5951** | ALS | 54 | Female | 7.2 | 12 |
| 5955 | ALS | 58 | Male | 7.1 | 3 |
| **5974** | ALS | 56 | Male | 7.9 | 22 |
| **6290** | ALS | 61 | Male | 8.2 | 11 |
| **6314** | ALS | 68 | Female | 7.4 | 7 |
| **6350** | ALS | 60 | Male | 8.3 | 10 |

**Table S2:** Target ALS cervical spinal cord samples for pRIPK1 and total RIPK1 assessment by MSD PMI: postmortem interval.

| Donor ID | Diagnosis | Age | Sex | PMI (h) | Family History of ALS/FTD |
| --- | --- | --- | --- | --- | --- |
| NEUCG551GV1 | Control | 57 | Female | 32 | Unknown |
| NEUEG442LDR | Control | 22 | Male | 6 | Unknown |
| NEUGJ583BEE | Control | 70 | Male | 91 | Unknown |
| NEUGY741GH8 | Control | 59 | Male | 18 | Unknown |
| NEUHA141EEC | Control | 72 | Male | 14 | Unknown |
| NEUHD481VCL | Control | 54 | Female | 7 | Unknown |
| NEUNJ406FRR | Control | 16 | Female | 9.5 | Unknown |
| NEUWD570BTK | Control | 68 | Male | 10 | Unknown |
| NEUXR099CKK | Control | 50 | Female | 5.5 | Unknown |
| NEUBB078VEG | ALS Spectrum MND | 51 | Female | 7 | Unknown |
| NEUCV649UJK | ALS Spectrum MND | 80 | Male | 5.3 | Unknown |
| NEUCW292DYJ | ALS Spectrum MND | 67 | Female | 7 | Unknown |
| NEUGV399KJW | ALS Spectrum MND | 63 | Female | 37.5 | Unknown |
| NEUGY575NUQ | ALS Spectrum MND | 69 | Male | 11.5 | Unknown |
| NEUMT947ALA | ALS Spectrum MND | 74 | Male | 3.5 | Unknown |
| NEURK749WFB | ALS Spectrum MND | 65 | Female | 2 | Yes |
| NEURX991BWB | ALS Spectrum MND | 59 | Female | 10 | No |
| NEUTP019MY9 | ALS Spectrum MND | 61 | Female | 5.5 | Unknown |
| NEUZB953VCD | ALS Spectrum MND | 68 | Female | 14 | Unknown |
| NEUEA633EN5 | ALS Spectrum MND | 71 | Female | 11 | Unknown |
| NEUZX073BDK | ALS Spectrum MND | 68 | Female | 6 | Unknown |
| JHU38 | ALS Spectrum MND | 34 | Female | 5 | Unknown |
| NEUEF397VVN | ALS + AD | 76 | Female | 8 | Unknown |
| NEUEH612JPA | ALS + AD | 71 | Female | 17.5 | Unknown |
| NEUKY135CV6 | ALS + AD | 74 | Male | 7 | Unknown |
| NEUNL548ZBN | ALS + FTD | 60 | Male | 12 | Yes |
| NEUUW911LWA | ALS + FTD | 59 | Male | 10 | Yes |
| NEUYX020CB0 | ALS + AD | 67 | Male | 7.5 | Unknown |

**Table S3:** Mass General Hospital spinal cord samples for pRIPK1 and total RIPK1 assessment by MSD RIN: RNA integrity number, PMI: postmortem interval.

| Donor ID | Diagnosis | Age | Sex | RIN | PMI (h) | Spinal cord region |
| --- | --- | --- | --- | --- | --- | --- |
| 2068 | Control | 79 | Female | 5.4 | 9 | Unknown |
| 2018 | Control | >=90 | Female | 4.6 | 24 | High cervical |
| 1887 | Control | 60 | Male | 6.3 | 14 | High cervical |
| 1805 | Control | 71 | Male | 2.25 | 72 | Unknown |
| 1721 | Control | 87 | Male | 3 | 24 | Thoracic |
| 1026 | Control | 69 | Male | N/A | 24 | Unknown |
| 2421 | ALS | 34 | Male | 5.05 | 26 | Cervical |
| 2418 | ALS | 62 | Female | 6.1 | 15 | Cervical |
| 2412 | ALS | 40 | Male | 7.85 | 42 | Cervical |
| 2402 | ALS | 51 | Male | 6.9 | 41 | Unknown |
| 2345 | ALS | 60 | Female | 5.65 | 31 | Lumbar |
| 2343 | ALS | 61 | Male | 6 | 16 | Cervical |
| 2322 | ALS | 60 | Male | 7.8 | 5 | Thoracic |
| 2319 | ALS | 60 | Male | 5.65 | 24 | Unknown |
| 2309 | ALS | 74 | Female | 5.2 | 48 | Thoracic |
| 2300 | ALS | 65 | Male | 6.15 | 96 | Thoracic |
| 2294 | ALS | 51 | Male | 6.85 | 24 | Thoracic |
| 2255 | ALS | 63 | Male | 7.6 | 30 | Unknown |
| 2253 | ALS | 66 | Male | 5.9 | 18 | Thoracic |
| 2251 | ALS | 70 | Female | 6.6 | 7 | Unknown |
| 2241 | ALS | 54 | Female | 5.05 | 20 | Thoracic |
| 2235 | ALS | 70 | Male | 5.75 | 26 | Thoracic |
| 2230 | ALS | 71 | Female | 7.1 | 30 | Thoracic |
| 2222 | ALS | 66 | Male | 5.9 | 19 | Unknown |
| 2216 | ALS | 52 | Male | 7.15 | 36 | Unknown |
| 2213 | ALS | 74 | Female | 8.4 | 18 | Unknown |
| 2117 | ALS | 61 | Male | 7.6 | 24 | Unknown |
| 2025 | ALS | 69 | Female | 7.4 | 36 | Unknown |

**Table S4:**
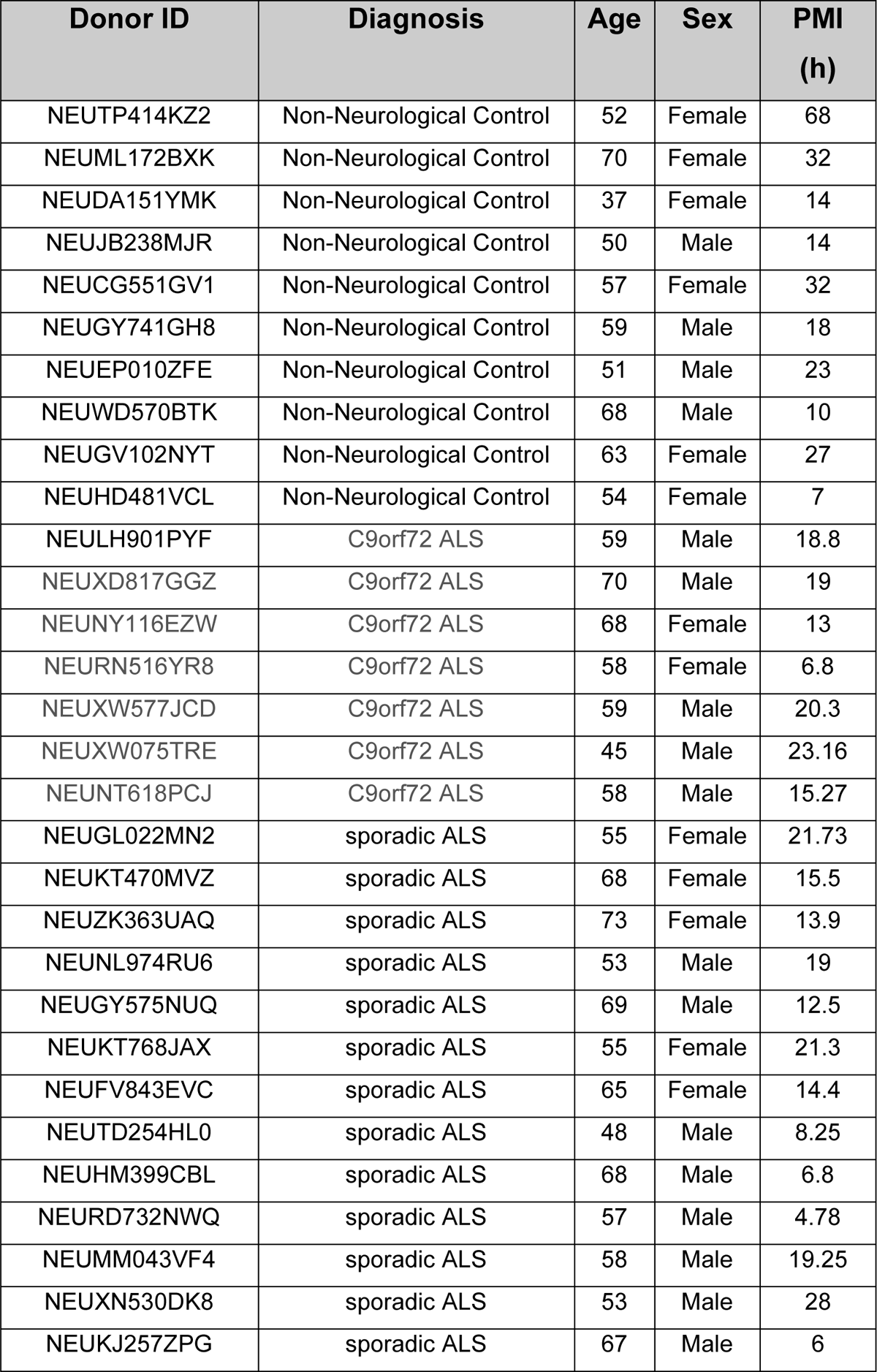
Target ALS thoracic spinal cord and motor cortex samples for total RIPK1 assessment by MSD RIN: RNA integrity number, PMI: postmortem interval.

**Table S5:** PrecisionMed CSF samples for NULISA cytokine analysis.

| Donor ID | Diagnosis | Age | Sex |
| --- | --- | --- | --- |
| 4830 | Definite ALS | 41 | Male |
| 4833 | Definite ALS | 48 | Male |
| 4850 | Definite ALS | 48 | Male |
| 4851 | Probable ALS | 48 | Male |
| 4816 | Definite ALS | 50 | Male |
| 4832 | Definite ALS | 57 | Male |
| 4825 | Definite ALS | 61 | Male |
| 4834 | Definite ALS | 67 | Male |
| 4820 | Definite ALS | 69 | Male |
| 4803 | Definite ALS | 70 | Male |
| 7550 | Control | 49 | Male |
| 7612 | Control | 56 | Male |
| 7600 | Control | 60 | Male |
| 7608 | Control | 65 | Male |
| 8202 | Control | 69 | Male |
| 7539 | Control | 43 | Male |
| 7519 | Control | 45 | Male |
| 7551 | Control | 49 | Male |
| 7557 | Control | 53 | Male |
| 7598 | Control | 71 | Male |

## Acknowledgements

Human tissue samples were obtained from the NIH NeuroBioBank (University of Pittsburgh, University of Maryland, and Massachusetts General Hospital) and through Target ALS. For the Target ALS data, we would like to thank the Target ALS Human Postmortem Tissue Core, the New York Genome Center for Genomics of Neurodegenerative Disease, the Amyotrophic Lateral Sclerosis Association and the TOW Foundation, Discovery Bioinformatics Services, and Qiagen Digital Insights (Qiagen Discovery Bioinformatics Services). Biofluid samples were obtained from PrecisionMed. Illustrations were created with BioRender. Research support funds were received from Denali Therapeutics Inc. and Sanofi.

## Author contributions

Conceptualization: M.Z., D.O., and T.R.H. Methodology: M.Z., D.O., and T.R.H. Investigation: M.Z., A.B., F.P., M.L., J.H., O.E.T., Y.R. S.K.R., P.K., M.L., D.S., Y.C., J.G., X.T., J.H., F.H., B.Z., G.G., D.O., and T.R.H. Visualization: M.Z., A.B., and T.R.H. Project administration: D.O. and T.R.H. Supervision: D.O. and T.R.H. Writing—original draft: M.Z.

## Materials and Methods

### Data availability

Target ALS datasets are available through https://www.targetals.org/resource/genomic-datasets/. Transcriptomic data will be deposited and accessible through GEO or Synapse (https://www.synapse.org/) by the time of publication.

### Human ALS bulk RNA sequencing

#### Tissue processing, library preparation and sequencing

Total RNA was purified from the spinal cord tissue homogenate using the RNeasy lipid tissue mini kit (Qiagen, 74804) with phase-lock tubes (Quantabio, 2302830) following manufacturer’s instructions. 100ng input RNA was used for library preparation with the NEBNext Globin & rRNA depletion kit (New England Biolabs, E7750X) according to manufacturer’s instructions, with 13 cycles for PCR. The samples were sequenced on an Illumina NovaSeq 6000.

#### Bulk RNA-seq analysis

Data were analyzed with Array Studio (Version 11, Omicsoft Corporation). Reads were mapped to the human B38 reference genome with the Ensembl R109 gene model using the Omicsoft Aligner 4 (OSA4) (54). Quantification was performed at gene level to obtain count-level information using the Expectation-Maximization (EM) algorithm (55). Genes were removed from the analysis if their counts equaled 0 for 9 or more samples. To identify transcriptomic differences between ALS samples and control samples, differential expression analysis was performed on the pseudobulk counts matrices while controlling for sex using DESeq2 version 1.38.3 in R version 4.2.2. Differentially expressed genes were identified using absolute(fold-change) >= 1.5 and FDR <= .05 criteria. Heatmaps of differentially expressed genes were plotted using the Pheatmap package version 1.0.12 in R using unbiased hierarchical clustering.

### Human ALS single nucleus RNA sequencing

#### Tissue processing, library preparation and sequencing

Human spinal cord samples were homogenized with a dounce homogenizer (Sigma Aldrich, D8938) in 1mL nuclei lysis buffer using the Nuclei Pure Prep Nuclei Isolation Kit (Sigma Adlrich, NUC-201). 3mL lysis buffer was added and incubate on ice for 15 minutes. 200µL of the total 4mL homogenate was separated for bulk RNA sequencing after adding 800uL Qiazol (Qiagen, 79306) and freezing samples at −80°C. Remaining homogenate was filtered through a 70µm strainer and 1mL cell suspension was layered on a 1.8M sucrose gradient and spun at 30,000g for 45 minutes at 4°C in a Beckman UltraCentrifuge. Cells were stained with DAPI at 20µg/mL (Life Technologies, D3571) and filtered through 40µm FlowMi cell strainers (Thermo Fisher Scientific, 14100150) prior to FACS sorting for DAPI+ nuclei on the Influx (BD Biosciences). Cells were loaded onto the 10x Genomics Next GEM Single Cell 3′ v3.1 sample chip and run following the manufacturer’s protocols. All 12 samples were processed in parallel and were amplified 12 cycles during the initial cDNA amplification and 12-13 cycles for the sample index PCR. Library quantification and quality assessment was performed with the Qubit 1x dsDNA High Sensitivity assay kit (Invitrogen, Q33231). The samples were sequenced on an Illumina NovaSeq 6000 to an average depth of approximately 50,000 reads per cell, from about 9,000 cells per sample. Data were combined from 2 independent sequencing runs for analysis.

#### Object setup, QC, and clustering

Samples were analyzed using Seurat 4.0.4 in R version 4.0.3. Samples were filtered to limit the inclusion of low-quality nuclei. Cells with fewer than 1,000 genes, 2,500 UMIs, and more than 15% mitochondrial genes were removed from analysis. After filtering, the dataset was normalized and the variable features were selected using the ‘vst’ method and using 2,000 features. The dataset was then scaled and RunPCA was performed for 100 principal components (PCs), and then the FindNeighbors function was run using 80 PCs to build the k-nearest neighbor (KNN) graph. Cluster resolution was set to ‘0.55’, and the uniform manifold approximation and projection (UMAP) was generated using the RunUMAP function and using 80 PCs. Cluster identities were determined by calculating enriched markers using the FindAllMarkers function in Seurat. Cell types were assigned by identifying genes unique to each cluster and by cross-referencing known markers of each cell type from existing published datasets. Doublet subpopulations were identified based on co-expression of core cell marker genes. These subpopulations were removed from the dataset, including a neuron-oligodendrocyte and astrocyte-microglia subpopulation. Following removal the dataset was reclustered using 75 PCs and a resolution of ‘0.55’. UMAP plots and gene expression plots were generated using built-in Seurat/ggplot2 plotting functions or scCustomize version 2.0.1 unless otherwise described.

#### Cell proportion quantification

To calculate the percent of cells per cluster from each sample, the number of cells from each sample in a given cluster was calculated and normalized to the number of cells per sample. These values were then normalized to the other replicate samples. Significantly enriched samples were identified using a two-way ANOVA with Tukey post hoc test, and P values are reported in each figure.

#### Pseudobulk analysis

Pseudobulk counts matrices were generated by summing the RNA counts for a given gene across all cells within a cell type for each individual sample. This was repeated for each cell type for which there was sufficient cell number (i.e., > 200 cells per condition) as well as for all cell types combined. To identify transcriptomic differences between ALS samples and control samples, differential expression analysis was performed on the pseudobulk counts matrices while controlling for sex using DESeq2 version 1.38.3 in R version 4.2.2. Differentially expressed genes were identified using absolute(fold-change) >= 1.5 and FDR <= .05 criteria. Heatmaps of differentially expressed genes were plotted using the Pheatmap package version 1.0.12 in R using unbiased hierarchical clustering.

#### Gene correlation analysis

The Pearson correlation coefficient and corresponding p-value were calculated between sample pRIPK1 levels and each gene using the pseudobulk counts matrices for either microglia or astrocytes.

#### Microglia and astrocyte subclustering and analysis

Microglia were subset from the larger dataset based on cell type definitions and reanalyzed using Seurat. The variable features were selected using the ‘vst’ method and using 2,000 features. RunPCA was performed for 20 PCs, and then the FindNeighbors function was run using 12 PCs to build the KNN graph. Cluster resolution was set to ‘0.1’, and the UMAP was generated using the RunUMAP function and using 12 PCs. Cluster/cell identities were determined by calculating enriched markers using the FindAllMarkers function in Seurat and small numbers of contaminating cells were removed based on expression of non-microglial markers. The dataset was then reclustered. RunPCA was performed for 20 PCs, and then the FindNeighbors function was run using 14 PCs to build the KNN graph. Cluster resolution was set to ‘0.15’, and the UMAP was generated using the RunUMAP function and using 14 PCs. UMAP plots and gene expression plots were generated using built-in Seurat/ggplot2 plotting functions unless otherwise described. To identify subcluster specific markers, each subpopulation was compared to the homeostatic cluster using the FindMarkers function and the “DESeq2” test with a minimum expression percentage of 0.25. Genes were rank ordered by adjusted p-value and log2 fold change. The percentage of cells per sample in each cluster was calculated by dividing the number of cells per sample in each cluster by the total number of cells in that sample. Significantly enriched samples were identified using a two-way ANOVA with Tukey post hoc test, and P values are reported in each figure.

Astrocytes were subset from the larger dataset based on cell type definitions and reanalyzed using Seurat. The variable features were selected using the ‘vst’ method and using 2,000 features. RunPCA was performed for 20 PCs, and then the FindNeighbors function was run using 12 PCs to build the KNN graph. Cluster resolution was set to ‘0.1’, and the UMAP was generated using the RunUMAP function and using 10 PCs. Cluster/cell identities were determined by calculating enriched markers using the FindAllMarkers function in Seurat but no contaminating cell types were found. UMAP plots and gene expression plots were generated using built-in Seurat/ggplot2 plotting functions unless otherwise described. To identify subcluster specific markers, each subpopulation was compared to the homeostatic cluster using the FindMarkers function and the “DESeq2” test with a minimum expression percentage of 0.25. Genes were rank ordered by adjusted p-value and log2 fold change. The percentage of cells per sample in each cluster was calculated by dividing the number of cells per sample in each cluster by the total number of cells in that sample. Significantly enriched samples were identified using a two-way ANOVA with Tukey post hoc test, and P values are reported in each figure.

#### Pathway and gene ontology analysis

ToppGene Suite (https://toppgene.cchmc.org/) and Ingenuity Pathway Analysis (QIAGEN) were used for gene ontology, pathway, and gene-disease association analysis on the mouse, human, and iPSC tri-culture datasets.

#### Target ALS analysis of RNA-seq data

Raw fastq RNA-Seq data files (1440 samples total) were provided by the NYGC ALS Consortium (Target ALS Release, July 2022; available through https://www.targetals.org/ resource/genomic-datasets/). These fastq files were generated using human postmortem tissue samples from the Target ALS postmortem tissue core and processed for RNA-Seq as previously described [62]. The raw fastq files were processed by Qiagen in OmicSoft ArrayStudio RNA-Seq analysis Pipeline (version 11.7.3.3). In brief, the fastq files had quality control performed and then aligned to the Genome Reference Consortium Human Build 38 using the proprietary OmicSoft Aligner [63]. After alignment, the gene level RPKM/FPKM/counts were determined using the EM algorithm in ArrayStudio as described previously [64]. Finally, pairwise differentially expressed genes were calculated using the DESeq2 v1.10.1 [65] in ArrayStudio comparing tissue-specific ALS patients versus non-neurological controls. ALS patients with multiple neurological conditions were excluded from pairwise analysis. Heatmaps were generated using Excel. Differentially expressed genes (DEGs) were considered significant with FDR < 0.05.

#### Human tissue processing and sample preparation

Fresh frozen spinal cord and motor cortex tissue was obtained through the NIH NeuroBioBank and from Target ALS (Tables S1 to S5). Cohorts included samples from the University of Maryland, University of Pittsburgh, and Massachusetts General Hospital. All lysis buffers contained protease and phosphatase inhibitors (Thermo Fisher Scientific, 78444) and tissue homogenization was carried out on ice. Post-mortem spinal cord tissue samples were homogenized and lysed in 1% Triton lysis buffer (TBS) at a 5x ratio of tissue weight to lysis buffer volume. Samples were centrifuged (13,000g) and the supernatant was collected as the soluble protein fraction. The pellet was further lysed with RIPA buffer (Boston BioProducts, BP-115) and sonicated with a fine tip sonicator to obtain the insoluble protein fraction. Tissue samples in Figures S5C and S5D were were obtained from TargetALS and homogenized in 6x tissue weight of 1x CST lysis buffer (Cell Signaling Technology, 9803) by TissueLyser. After homogenization, samples were incubated on ice for 30 minutes and then centrifuged at max speed for 30 minutes. The supernatant was transferred to new tubes and used as the soluble fraction. Protein concentration for all samples was determined by BCA assay.

#### RIPK1 MSD ELISA

Biotinylated RIPK1 capture antibody (BD Bioscience #610459) was diluted in PBS to 1mg/mL and added to MSD Gold 96-well small spot streptavidin plates and incubated with shaking (700rpm) for 1 hour at RT. After blocking with MSD Blocker A for 2 hours with shaking (700rpm), 25-30 µL of tissue lysates were incubated overnight at 4°C with shaking (600rpm). For human tissue phospho-RIPK1 (Cell Signaling Technology, 65746) or total RIPK1 antibodies (Cell Signaling Technology, 3493) were diluted 1:500 in MSD Blocker A and added as detection antibodies for 1.5 hours with shaking (700rpm) at RT.

Secondary detection antibody (goat anti-rabbit antibody, sulfo-TAG labeled, Meso Scale Discovery, R32AB-1) was diluted 1:1000 in MSD Blocker A and added for 1 hour at RT with shaking (700rpm). For target engagement assays through detection of free and total RIPK1 as described in [50], mouse blood was lysed 1:1 with 2x CST lysis buffer with protease and phosphatase inhibitor cocktail. Free and total RIPK1 levels were separately detected with total RIPK1 antibodies (Cell Signaling Technology, 3493 and abcam, ab202985). Human CSF samples were not lysed and free and total RIPK1 levels were separately detected with (Cell Signaling Technology, 3493 and abcam, ab125072). The assay was developed with 150 mL of 2x MSD read buffer (R92TC-3) and plates were read on an MSD Meso Sector imager S600. ECL counts were normalized to protein concentration after blank subtraction. For target engagement the ratio of free to total RIPK1 was normalized to the baseline to calculate percent change relative to baseline.

#### Transgenic mice

All animal studies were conducted in compliance with the ethical regulations and full approval of Sanofi’s Institutional Animal Care and Use Committee (IACUC). Female B6.Cg-Tg(SOD1*G93A)1Gur/J (Jackson Laboratory strain 004435) mice and non-transgenic C57BL6/J controls were obtained from the Jackson Laboratory at 60 days of age. The mice were maintained in a humidity- and temperature-controlled vivarium (20– 22°C) on a 12/12 h light/dark schedule. Animals had access ad libitum to food and water. Mice were dosed with RIPK1 inhibitor (RIPK1i) in chow at 100mg/kg starting on postnatal day 70 (Research Diets Inc., Purina Rodent Chow #5001 with 0.667g RIPK1i/kg diet (C21020201i)). On postnatal day 135 twice daily dosing (BID) of vehicle (0.5% methyl cellulose in water) or RIPK1i (60mg/kg) was initiated [29]. Symptom onset and disease progression were monitored as described [66], with abnormal splay or collapse of hindlimb monitored for symptom onset. Mice were assessed 3 times a week for body weight change and symptom onset or progression. Mice were euthanized at clinical endpoint defined by either 20% weight loss from peak body weight or complete hindlimb paralysis.

#### Behavioural analyses

For the wire hang test, mice were suspended from a wire mesh for a maximum of 180 seconds, and the latency to fall was measured for each individual mouse. Mice were assessed every 10 days from postnatal day 90 to postnatal day 140. For the open field test to assess locomotor activity, individual mice were placed in a Plexiglass cage and allowed to explore for a few minutes (San Diego Instruments, Photobeam Activity System. Computer-aided video tracking system (Photobeam Activity System, Version 2) was used to measure the distance traveled, speed of movement, and rearing on hindlimbs in a 20 minute period. A photobeam array placed around the exterior of the cage recorded each animal’s movement.

#### Plasma Neurofilament-H

Murine plasma neurofilament-heavy (NF-H) was assessed on 5μL murine plasma using the human SimplePlex Assay as the kit is cross reactive with murine NF-H (R&D Systems, Protein Simple SPCKB-PB-000519) according to manufacturer’s instructions. The SimplePlex Assay performed on the Ella platform utilizes a microchip with capillaries coated with NF-H antibodies.

### SOD1^G93A^ mouse spinal cord single cell RNA sequencing

#### Mouse spinal cord processing and library preparation

At 120 days of age, C57BL/6 mice along with vehicle and RIPK1i-treated SOD1^G93A^ mice were sacrificed and perfused with HBSS supplemented with 5 µg/mL actinomycin D and 10 µM triptolide. The spinal cords were isolated, cut sagitally, and placed in 10mL pre-incubation solution (HBSS (Gibco, 14025-092) supplemented with transcription/translation inhibitor cocktail: 5 µg/mL actinomycin D, 10 µM triptolide, and 27.1 µg/mL anisomycin) on ice and protected from light. Spinal cords were minced on ice with a razor blade then digested for 15 minutes on a shaker at 37°C in 5mL papain solution (300U papain solution (Worthington, LS003126) was diluted in HBSS supplemented with 1:100 DNAse and the transcription/translation inhibitor cocktail). Every 5 minutes samples were triturated with a glass pipette and the papain reaction was quenched with 5mL HBSS supplemented with 10% FBS. After filtering through a 70µm strainer and pelleting cells for 5 minutes at 300g and 4°C, samples were resuspended in 3mL HBSS with 10% FBS and 7mL of 25% BSA solution in HBSS supplemented with the transcription and translation inhibitor cocktail. Cells were centrifuged for 10 minutes at 1200g and the supernatant and myelin were aspirated prior to addition of 1mL cold HBSS and a 2 minute spin at 500g. Finally, the solution was decanted and cells were quantified after resuspending in 500µL HBSS. Cells were loaded onto the 10x Genomics Next GEM Single Cell 3′ v3.1 sample chip and run following the manufacturer’s protocols. All 12 samples were processed in parallel and were amplified 12 cycles during the initial cDNA amplification and 14 cycles for the sample index PCR before sequencing on an Illumina NovaSeq 6000.

#### Object setup, QC, and clustering

Samples were analyzed using Seurat 5.0.1 in R version 4.2.2. Samples were filtered to limit the inclusion of low-quality cells. Cells with fewer than 1,450 genes, more than 7,000 genes, and more than 5% mitochondrial genes were removed from analysis. After filtering, the dataset was normalized and the variable features were selected using the ‘vst’ method and using 2,000 features. The dataset was then scaled and RunPCA was performed for 65 principal components (PCs), and then the FindNeighbors function was run using 30 PCs to build the k-nearest neighbor (KNN) graph. Cluster resolution was set to ‘0.3’, and the t-distributed Stochastic Neighbor Embedding (TSNE) was generated using the RunTSNE function and using 30 PCs. Cluster identities were determined by calculating enriched markers using the FindAllMarkers function in Seurat. Cell types were assigned by identifying genes unique to each cluster and by cross-referencing known markers of each cell type from existing published datasets. TSNE plots and gene expression plots were generated using built-in Seurat/ggplot2 plotting functions or scCustomize version 2.0.1 unless otherwise described.

#### Pseudobulk analysis

Pseudobulk counts matrices were generated by summing the RNA counts for a given gene across all cells within a cell type for each individual sample. This was repeated for each cell type for which there was sufficient cell number (i.e., > 200 cells per condition) as well as for all cell types combined. To identify transcriptomic differences between conditions, differential expression analysis was performed on the pseudobulk counts matrices using DESeq2 version 1.38.3 in R version 4.2.2. Differentially expressed genes were identified using absolute(fold-change) >= 1.5 and unadjusted p-value <= .05 criteria.

### Human iPSC tri-culture single cell RNA sequencing

#### Tri-culture plating and stimulation

Tri-culture was plated as described previously in Ryan et al [18]. In brief, 96-well plates (Corning, 3595) were coated with Matrigel (Corning, 354277) on day 0 and iAstrocytes (Fujifilm, ASC-100-020-001-PT) were thawed and plated at 1.5 x 10^4^ cells/well in 200µL Fuji-designated astrocyte media (DMEM/F12, HEPES (Life Technologies, 11330057) with 2% heat-inactivated FBS Certified One Shot (Gibco, A38400-01) and 1x N-2 supplement (Gibco, 17502-048)). On day 1, astrocyte media was fully aspirated and 3.5 x 10^4^ iCell motor neurons (Fujifilm, C1048) were added per well in 200µL complete Fuji motor neuron media (100 mL iCell Neural Base Medium 1 (Fujifilm, M1010) with 2 mL iCell Neural Supplement A (Fujifilm, M1032) and 1 mL iCell Nervous System Supplement (Fujifilm, M1031)). On day 5, iCell Microglia (Fujifilm, C1110) were plated at 1 x 10^4^ cells/well in complete tri-culture media (50mL motor neuron media supplemented with 0.5mL iCell Microglia Supplement A (Fujifilm, M1036), 0.5mL iCell Microglia Supplement B (Fujifilm, M1037) and 2mL iCell Neural Supplement C (Fujifilm, M1035)). Half media exchanges were performed every 2 to 3 days, with treatments added on day 15. Cells were pre-treated for 1 hour with 1µM Smac mimetic LCL161 (Selleck Chem, S7009), 20µM zVAD-fmk (Enzo Life Sciences, ALX-260-020), and 1µM RIPK1 inhibitor [29] prior to a 4 hour incubation with 50ng/mL recombinant human TNF (R&D Systems, 210-TA).

#### Tri-culture sample processing and library preparation

Four hours after treatment, cells were washed twice with PBS then treated with 0.25% trypsin-EDTA solution (Sigma-Aldrich, T4049) and transcription/translation inhibitors: 5 µg/mL actinomycin D (Sigma-Aldrich, A1410), 10 µM triptolide (Sigma-Aldrich, T3652), and 27.1 µg/mL anisomycin (Sigma-Aldrich, A9789) for 6 minutes at 37°C in a 5% CO_2_ incubator. Then, 1 volume of PBS with 2% FBS (Gibco, A38400-01), the transcription/translation inhibitors, and 1:100 DNAse (Worthington, LS002007) was added to quench. Cells from two wells were combined for each technical replicate and put through 40-µm cell strainers (BD Falcon, 352235). Cells were counted and spun at 4 °C for 5 minutes at 1500 rpm. Supernatant was removed and cells were resuspended at 1,000 cells/µL in ice-cold PBS. Cells were loaded onto the 10x Genomics Next GEM Single Cell 3′ v3.1 sample chip and run following the manufacturer’s protocols. All 10 samples were processed in parallel and were amplified 11 cycles during the initial cDNA amplification and 12 cycles for the sample index PCR before sequencing on an Illumina NovaSeq 6000.

#### Object setup, QC, and clustering

Samples were analyzed using Seurat 5.0.1 in R version 4.2.2. Samples were filtered to limit the inclusion of low-quality cells. Cells with fewer than 1,000 genes, more than 9,000 genes, or more than 15% mitochondrial genes were removed from analysis. After filtering, the dataset was normalized and the variable features were selected using the ‘vst’ method and using 2,000 features. The dataset was then scaled and RunPCA was performed for 100 principal components (PCs), and then the FindNeighbors function was run using 75 PCs to build the k-nearest neighbor (KNN) graph. Cluster resolution was set to ‘0.3’, and the uniform manifold approximation and projection (UMAP) was generated using the RunUMAP function and using 75 PCs. Cluster identities were determined by calculating enriched markers using the FindAllMarkers function in Seurat. Cell types were assigned by identifying genes unique to each cluster and by cross-referencing known markers of each cell type from existing published datasets. Doublet subpopulations were identified based on co-expression of core cell marker genes and were removed from the dataset. Following removal, the dataset was reclustered using 75 PCs and a resolution of ‘0.3’. UMAP plots and gene expression plots were generated using built-in Seurat/ggplot2 plotting functions or scCustomize version 2.0.1 unless otherwise described.

#### Pseudobulk analysis

Pseudobulk counts matrices were generated by summing the RNA counts for a given gene across all cells within a cell type for each individual sample. This was repeated for each cell type for which there was sufficient cell number (i.e., > 200 cells per condition) as well as for all cell types combined. To identify transcriptomic differences between conditions, differential expression analysis was performed on the pseudobulk counts matrices using DESeq2 (version 1.38.3) in R (version 4.2.2). Differentially expressed genes were identified using absolute(fold-change) >= 1.5 and unadjusted p-value <= .05 criteria. Heatmaps of differentially expressed genes were plotted using the Pheatmap package version 1.0.12 in R using unbiased hierarchical clustering.

#### Tri-culture stimulation for supernatant collection

Tri-culture was plated as described previously, but on day 14 media was exchanged twice at 75% with Neurobasal/B27+ media (Life Technologies, A3653401) supplemented with microglia growth factors: 25ng/mL M-CSF (PeproTech, 300-25), 100ng/mL IL-34 (PeproTech, 200-34), and 50ng/mL TGF-β1 (PeproTech, 100-21). On day 15 cells were pre-treated for 1 hour with 1µM Smac mimetic LCL161 (Selleck Chem, S7009), 20µM zVAD-fmk (Enzo Life Sciences, ALX-260-020), and 1µM RIPK1 inhibitor [29] prior to a 24 hour incubation with 50ng/mL recombinant human TNF (R&D Systems, 210-TA).

#### Olink analysis tri-culture supernatant

Proteomic analysis was conducted on the tri-culture supernatant with the Olink Explore Inflammation and Neurology panels. Raw data were analyzed using MyData software from Olink. After data normalization, protein concentration was expressed in Olink’s arbitrary unit in Log2 scale as NPX (Normalized Protein eXpression) values. Fold change of treated samples relative to controls was calculated as a difference of NPX_treated_-NPX_control_, where 1 unit of difference corresponds to a 2-fold difference in concentration. Cut-offs of >0.5 difference in NPX and p<0.05 were used for 2 separate comparisons: DMSO control vs TSZ, and TSZ vs TSZ + RIPK1i treatments. Statistical significance was established using a one-tailed t test.

#### Human CSF cytokine analysis with NULISA

Control and CSF samples were obtained from PrecisionMed. NULISAseq assays were performed at Alamar Biosciences as described previously [67]. Briefly, CSF samples stored at −80°C were thawed on ice and centrifuged at 10,000g for 10 minutes. 10µL supernatant samples were plated in 96-well plates and analyzed with Alamar’s Inflammation Panel 250, mainly targeting inflammation and immune response-related cytokines and chemokines. A Hamilton-based automation instrument was used to perform the NULISAseq workflow, starting with immunocomplex formation with DNA-barcoded capture and detection antibodies, followed by capturing and washing the immunocomplexes on paramagnetic oligo-dT beads, then releasing the immunocomplexes into a low-salt buffer, which were captured and washed on streptavidin beads. Finally, the proximal ends of the DNA strands on each immunocomplex were ligated to generate a DNA reporter molecule containing both target-specific and sample-specific barcodes. DNA reporter molecules were pooled and amplified by PCR, purified and sequenced on Illumina NextSeq 2000. Sequencing data were processed using the NULISAseq algorithm (Alamar Biosciences). The sample-(SMI) and target-specific (TMI) barcodes were quantified, and up to two mismatching bases or one indel and one mismatch were allowed. Intraplate normalization was performed by dividing the target counts for each sample well by that well’s internal control counts. Interplate normalization was then performed using interplate control (IPC) normalization, wherein counts were divided by target-specific medians of the three IPC wells on that plate. Data were then rescaled, add 1 and log2 transformed to obtain NULISA Protein Quantification (NPQ) units for downstream statistical analysis. CSF samples were run in duplicate so NPQ values for technical replicates were averaged. Unsupervised clustering using heatmaps and principal component analysis identified two outlier samples which were excluded. A linear regression model was fit to assess the difference in protein abundance between ALS and non-neurological control samples, where NPQ was modeled as the outcome and disease status was a binary predictor. Analyses were performed using R version 4.2.3. Statistical significance was defined using an unpaired two-tailed t test.

#### Human CSF cytokine analysis from Phase 1b trial with SAR443060

CSF samples from people with ALS obtained from the Phase 1b clinical trial (NCT03757351) with SAR443060 [49] were analyzed for cytokines. CCL2 levels in CSF diluted 20x with Diluent 43 were measured using the MSD V-Plex kit (# K151NND). CCL4 levels in CSF diluted 2x with Diluent 43 were measured using the MSD V-Plex kit (# K151NRD). Raw instrument values were interpolated against a calibration curve provided with the assay kit. Technical replicates for each sample were averaged and calculated as a percent change normalized to the baseline value for each patient.

#### Quantification and statistical analysis

All quantification and statistical analyses were completed in GraphPad Prism version 9.1.2, Array Studio Version 11, and R version 4.2.2 as described in the figure legends and methods. For tri-culture experiments 2-3 technical replicates as combined individual wells were used for sequencing and supernatant analysis. Biological replicates of individual mouse or human biofluid or tissue samples were used for all other experiments. Data are presented as mean ± SEM for all samples. For all panels where statistical significance is indicated, *P < 0.05, **P < 0.01, **P < 0.001 and ****P < 0.0001. For sequencing experiments, fold change cutoffs of |1.5| were used, with p < 0.05 for tri-culture and mouse spinal cords and FDR < 0.05 for human spinal cord samples. Statistical analysis across conditions was measured using unpaired one-tailed t test, unpaired two-tailed t test with Welch’s correction, log-rank test (Mantel-Cox), Gehan-Breslow-Wilcoxon test, one-way ANOVA with post hoc Dunn’s, or two-way ANOVA with post hoc Sidak’s multiple comparisons tests were used as indicated in the figure legends and methods.

