## Supplementary Figures for "Glial state changes and neuroinflammatory RIPK1 signaling are a key feature of ALS pathogenesis"

### Figure S1:

**A**

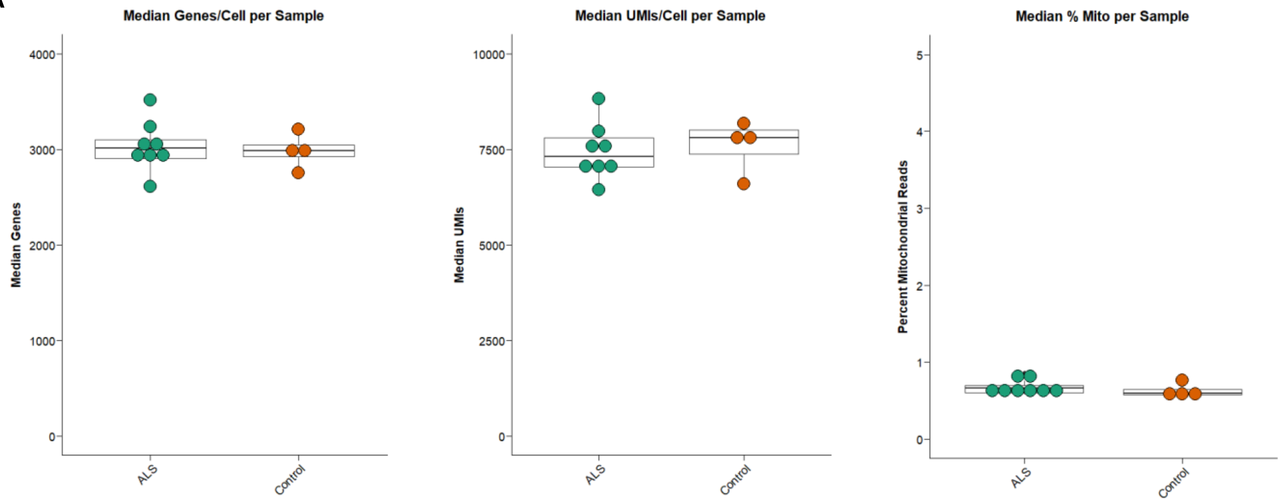


**B**

| Cell Type | % total cells |
|---|---|
| Oligodendrocyte | 51.70 |
| Microglia | 16.42 |
| Astrocyte | 14.40 |
| OPC | 6.81 |
| Fibroblast | 2.72 |
| Lymphocyte | 2.03 |
| Neuron | 2.00 |
| Endothelial | 1.42 |
| Pericyte/SMC | 0.79 |
| PVM | 0.73 |
| Ependymal | 0.72 |
| Schwann Cell | 0.26 |

**C**

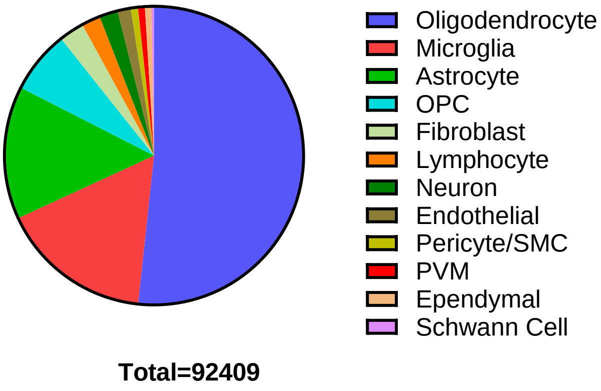


**D**

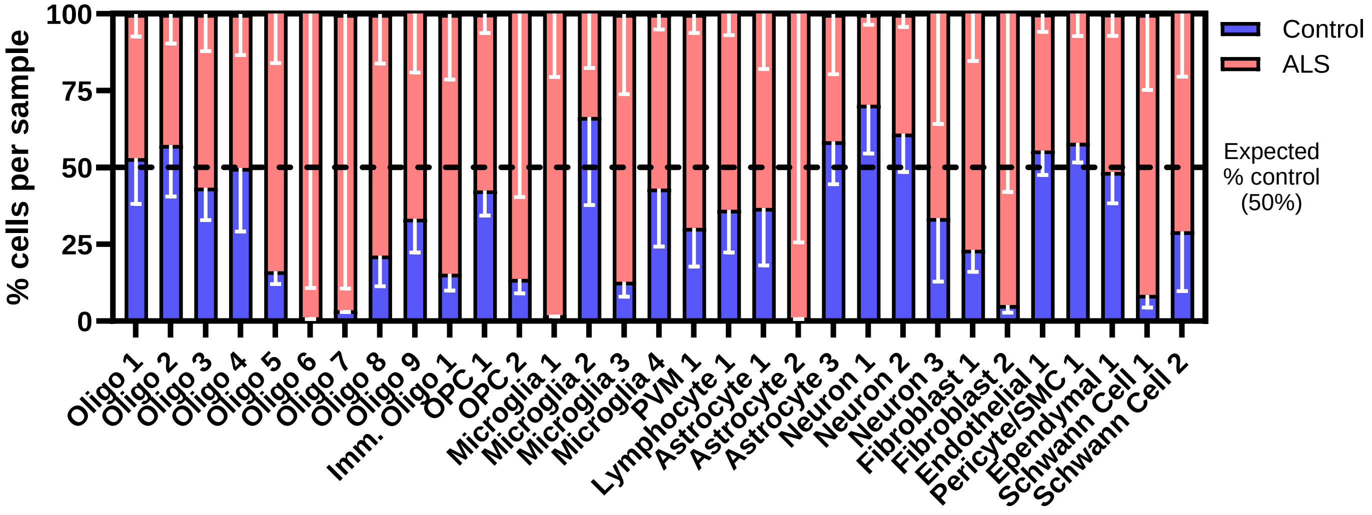

**Figure S1: Cell type distribution from snRNA-seq analysis of human ALS and control spinal cords**

A) Bar graphs depicting the median number of genes, unique molecular identifiers, and percent mitochondrial genes from snRNA-seq analysis of ALS (n=8) and control (n=4) spinal cords

B, C) Cell type distribution by percent (B) and number of total cells (C) from snRNA-seq analysis of ALS and control spinal cords

D) Bar graph depicting normalized percent of cells from each sample assigned to each subcluster from snRNA-sequenced ALS and control spinal cords

Data depict biological replicates and error bars represent mean ± SEM.

### Figure S2:

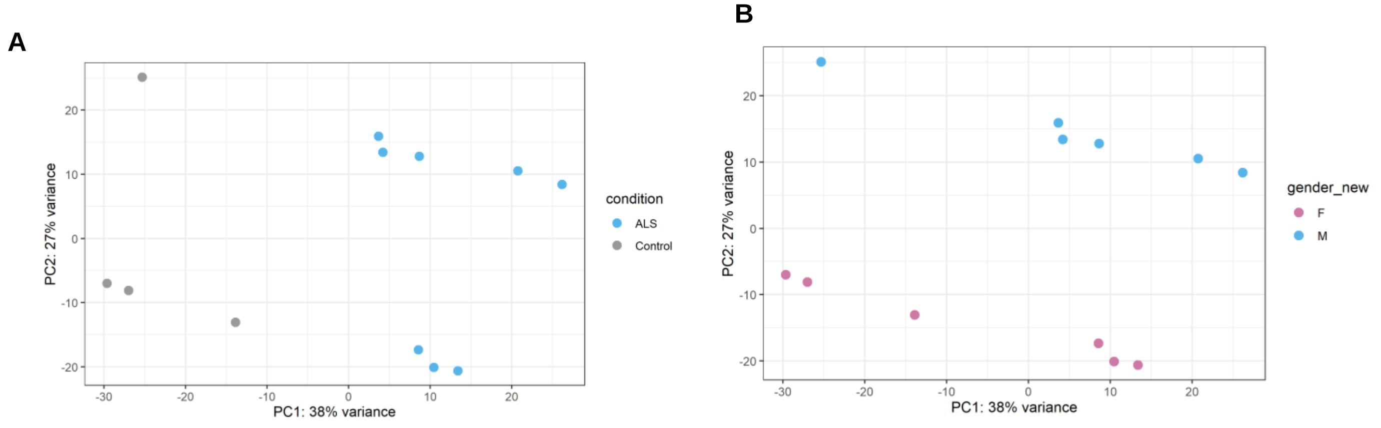


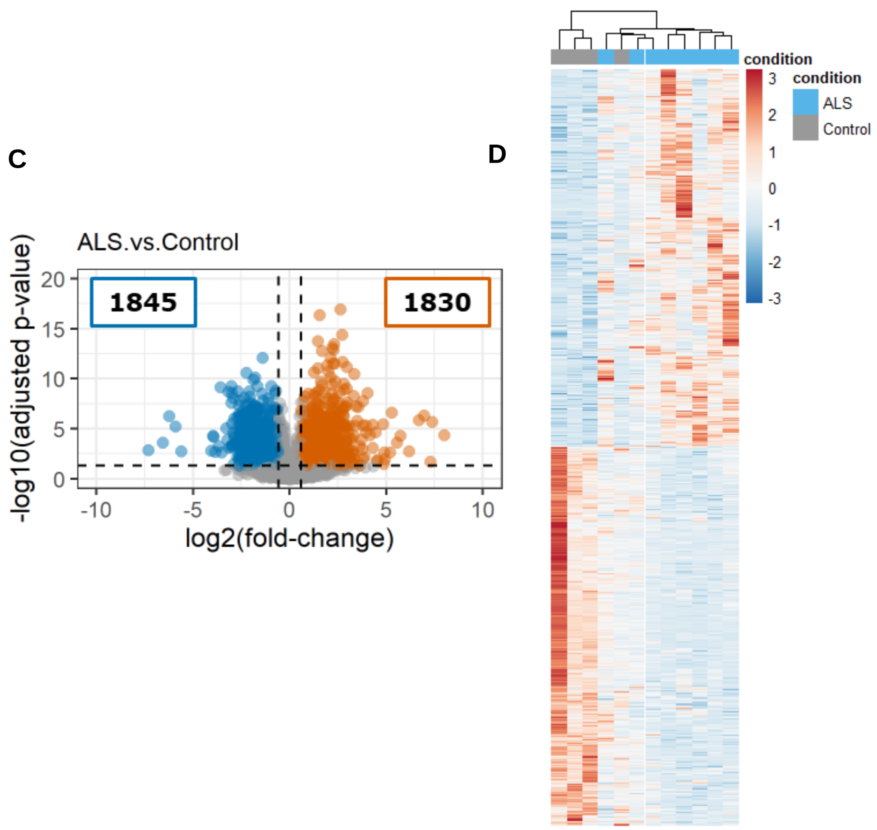


E

##### Upregulated DEGs

| Biological Process | pValue |
|---|---|
| regulation of immune system process | 2.09E-51 |
| cell activation | 1.22E-41 |
| inflammatory response | 2.96E-41 |
| leukocyte activation | 3.06E-37 |
| cytokine production | 3.55E-26 |

##### Downregulated DEGs

| Biological Process | pValue |
|---|---|
| synaptic signaling | 2.87E-87 |
| anterograde trans-synaptic signaling | 1.02E-85 |
| chemical synaptic transmission | 1.02E-85 |
| synapse organization | 3.20E-37 |
| neurotransmitter transport | 6.82E-28 |

**Figure S2: Characterization of genes and pathways modulated in ALS spinal cords from snRNA-seq analysis**

A, B) PCA of ALS (n=8) and control (n=4) spinal cords from snRNA-seq analysis grouped by disease (A) or gender (B)

C) Volcano plot showing DEGs in ALS versus control spinal cords analyzed by snRNA-seq (cutoff |FC| > 1.5, FDR < 0.05)

D) Hierarchical heatmap clustering of DEGs in ALS spinal cord samples relative to control from snRNA-seq pseudobulk analysis

E) Gene Ontology terms for biological processes from ToppGene Suite for upregulated and downregulated DEGs in ALS spinal cords relative to controls

Data depict biological replicates. FDR (Benjamini-Hochberg) was used (E).

### Figure S3:

A

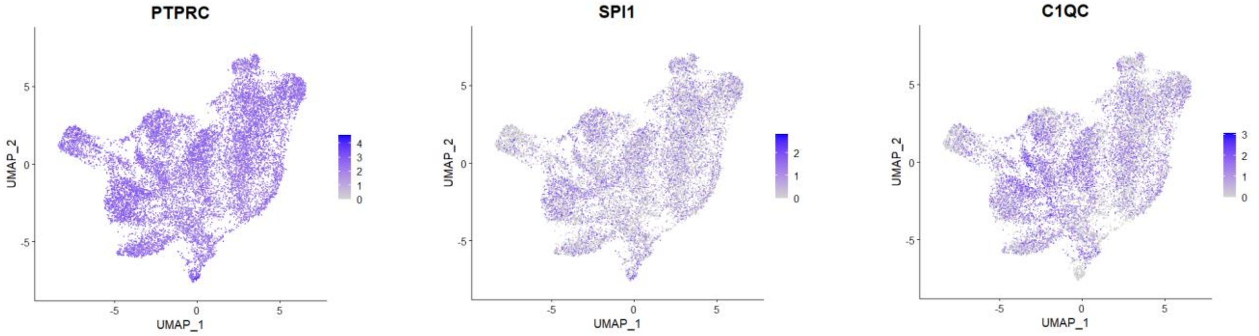


B

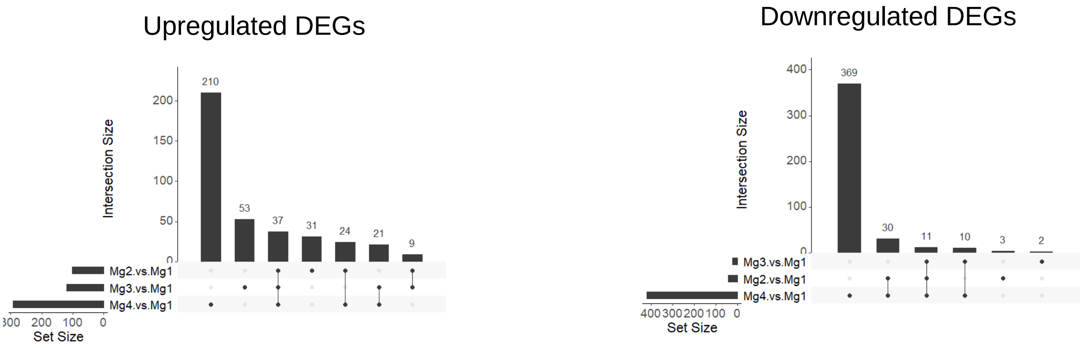


C

Mg 2

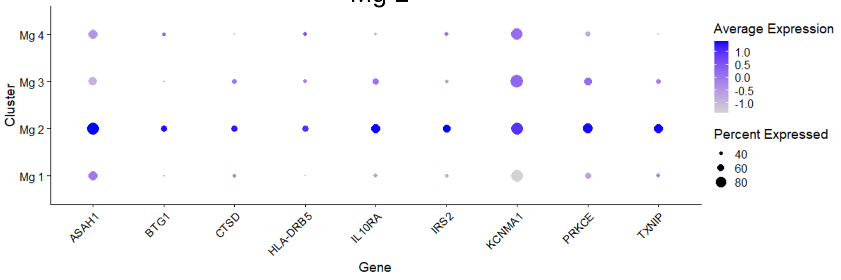


Mg 3

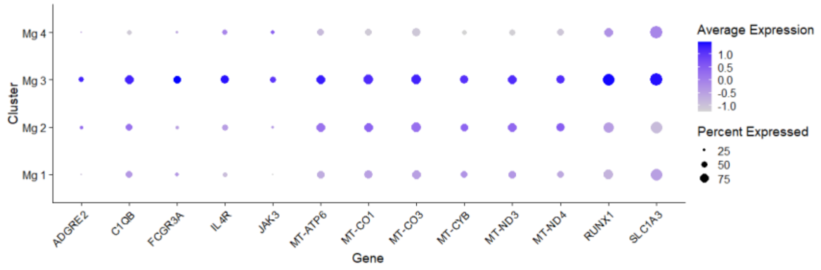


Mg 4

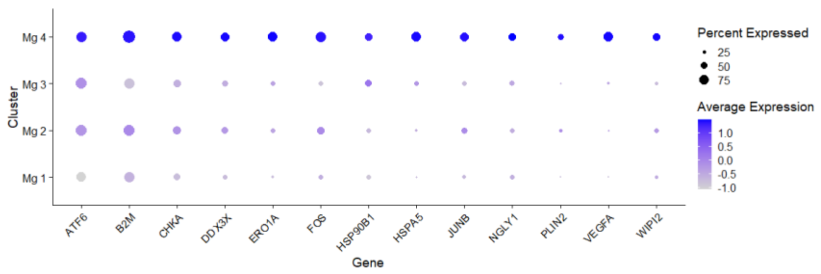


D

Mg 2

| Biological Process | pValue |
|---|---|
| regulation of lipid metabolic process | 3.35E-02 |
| response to hormone | 4.72E-02 |

Mg 3

| Biological Process | pValue |
|---|---|
| respiratory electron transport chain | 5.75E-04 |
| oxidative phosphorylation | 1.14E-03 |
| regulation of immune system process | 2.11E-03 |

Mg 4

| Biological Process | pValue |
|---|---|
| protein folding | 4.78E-04 |
| cellular response to starvation | 1.02E-03 |
| response to endoplasmic reticulum stress | 1.66E-03 |

**Figure S3: Characterization of microglia subclusters and ALS-related genes**

A) UMAP projections depicting microglial marker genes from snRNA-seq analysis of ALS (n=8) and control (n=4) spinal cords

B) Bar graph depicting the overlap of upregulated and downregulated DEGs across the microglial subclusters Mg 2 to Mg 4 relative to the homeostatic cluster Mg 1

C) Dot plots depicting unique upregulated DEGs in microglial subclusters Mg 2 to Mg 4 (cutoff |FC| > 1.5, FDR < 0.05)

D) Gene Ontology terms for biological processes from ToppGene Suite for unique upregulated DEGs in microglial subclusters Mg 2 to Mg 4

Data depict biological replicates. FDR (Benjamini-Hochberg) was used (D).

Internal

### Figure S4:

**A**

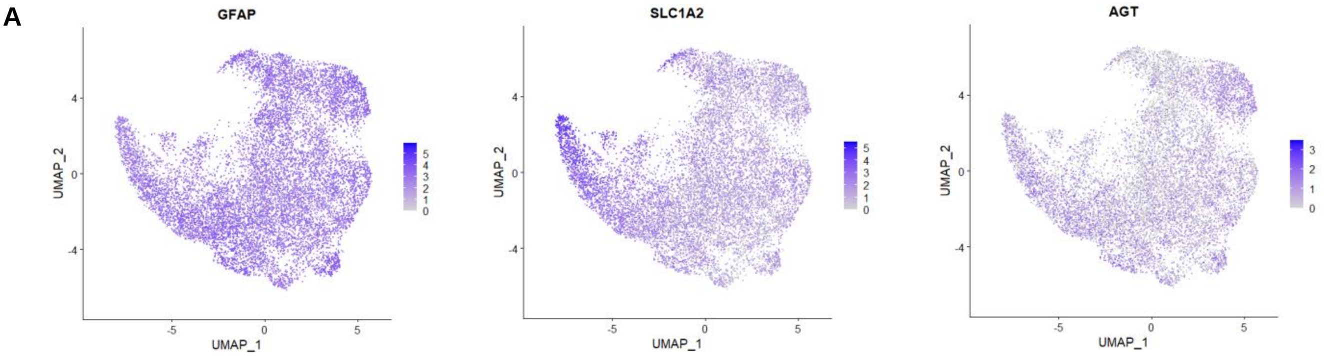


**B**

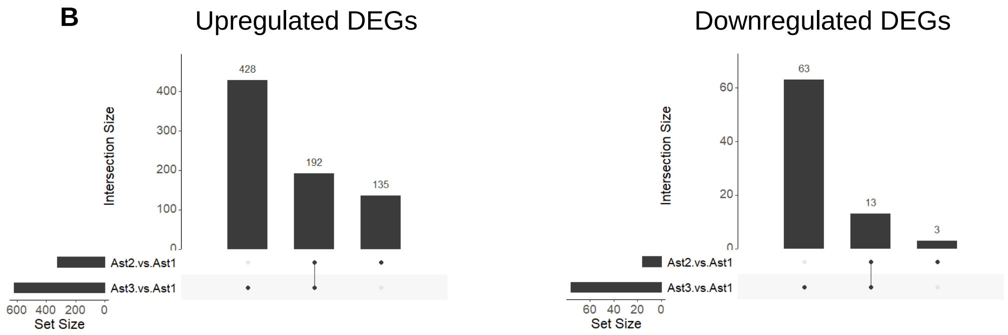


**C**

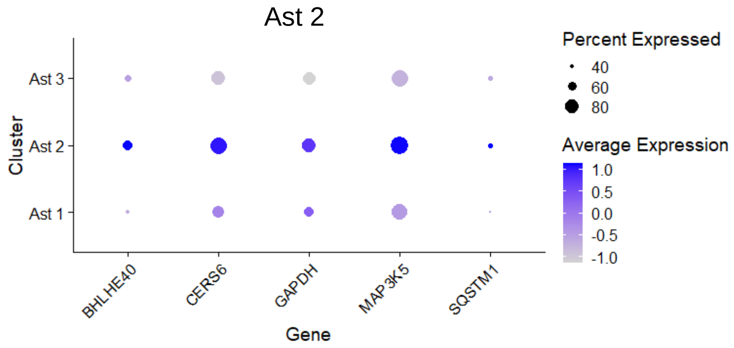


**D**

##### Ast 3 upregulated DEGs

| Disease | pValue | Source |
|---|---|---|
| Amyotrophic Lateral Sclerosis | 2.68E-10 | DisGeNET BeFree |
| Astrocytoma | 7.10E-07 | DisGeNET BeFree |
| Arteriosclerosis | 8.26E-07 | DisGeNET BeFree |
| Parkinson Disease | 8.55E-07 | DisGeNET BeFree |
| Diabetic Nephropathy | 9.61E-07 | DisGeNET BeFree |

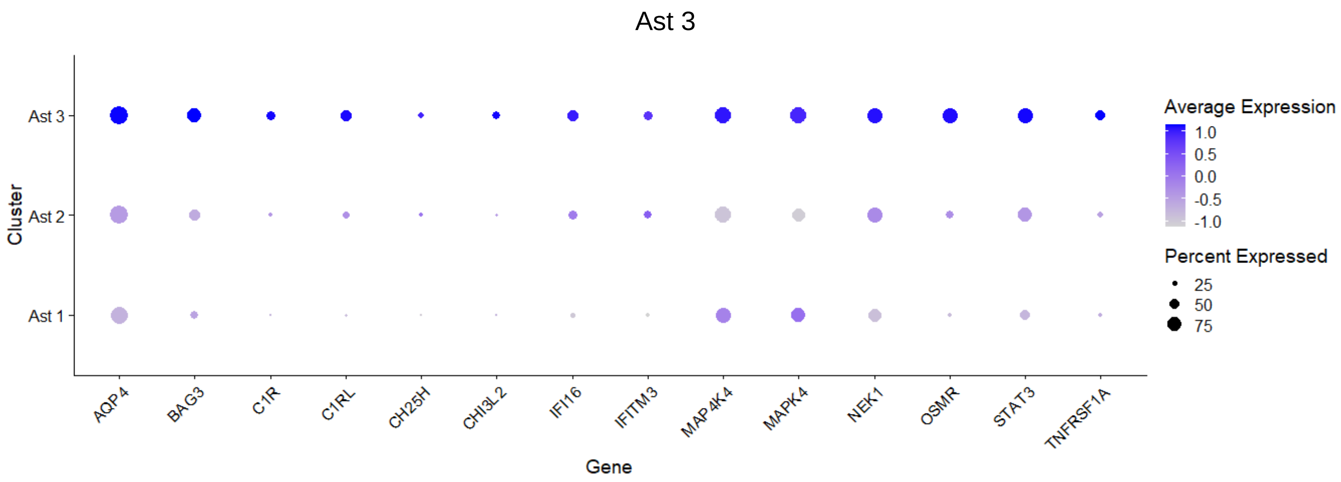

**Figure S4: Characterization of astrocyte subclusters and ALS-related genes**

A) UMAP projections depicting astrocyte marker genes from snRNA-seq analysis of ALS (n=8) and control (n=4) spinal cords

B) Bar graph depicting the overlap of upregulated and downregulated DEGs across the astrocyte subclusters Ast 2 and Ast 3 relative to the homeostatic cluster Ast 1

C) Dot plots depicting unique upregulated DEGs in astrocyte subclusters Ast 2 and Ast 3 (cutoff |FC| > 1.5, FDR < 0.05)

D) Most significant diseases for gene disease association from ToppGene Suite for upregulated Ast 3 DEGs

Data depict biological replicates. FDR (Benjamini-Hochberg) was used (D).

### Figure S5:

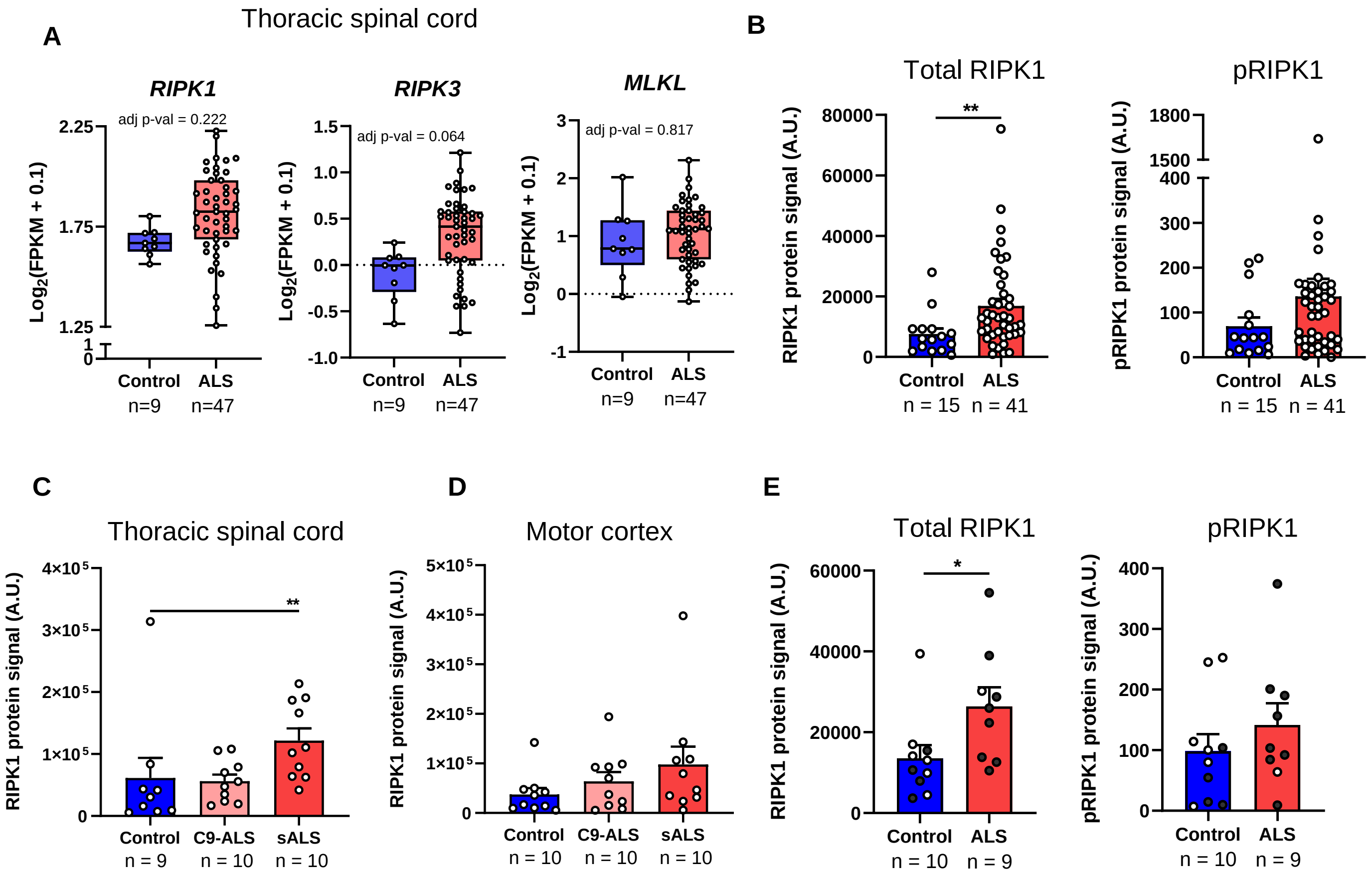
A
Thoracic spinal cord
RIPK1
RIPK3
MLKL
adj p-val = 0.222
adj p-val = 0.064
adj p-val = 0.817
Log2(FPKM + 0.1)
Control
ALS
n=9
n=47
B
Total RIPK1
pRIPK1
RIPK1 protein signal (A.U.)
pRIPK1 protein signal (A.U.)
**
n = 15
n = 41
C
Thoracic spinal cord
D
Motor cortex
C9-ALS
sALS
n = 9
n = 10
E
*

**Figure S5: RIPK1 expression and kinase activation in ALS and control postmortem tissue samples**

A) Gene expression levels of *RIPK1*, *RIPK3*, and *MLKL* in control and ALS thoracic spinal cord samples from the Target ALS bulk RNA-seq database

B) MSD assay quantifying total and pRIPK1 levels in the soluble (TBS-Triton) protein fraction from ALS (n=41) and control (n=15) spinal cords from Mass General Hospital and Target ALS cohorts

C, D) MSD assay quantifying total and pRIPK1 levels in soluble (CST lysis buffer) protein fraction from C9orf72 ALS, sporadic ALS, and control thoracic spinal cord (C) or motor cortex (D) samples from Target ALS (n=9-10 per group)

E) MSD assay quantifying total and pRIPK1 levels in soluble (TBS-Triton) protein lysates from ALS (n=9) and control (n=10) cervical spinal cords samples. Black dots depict samples used for bulk and single nucleus RNA sequencing

Data depict biological replicates and error bars represent mean ± SEM. FDR (A), unpaired two-tailed t test with Welch's correction (B, E) and one-way ANOVA (Kruskal-Wallis) with Dunn's multiple comparisons test (C) were performed. *$p < 0.05$, **$p < 0.01$; C9-ALS, C9orf72 ALS; sALS, sporadic ALS

### Figure S6:

#### Microglia

A

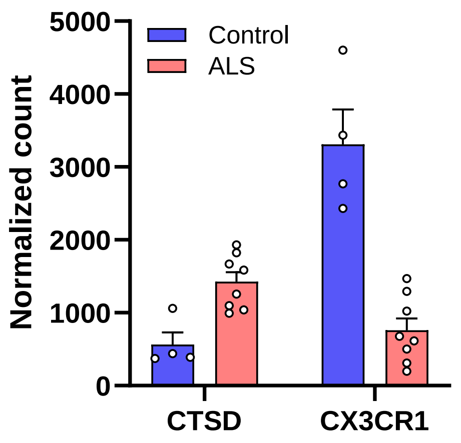


B

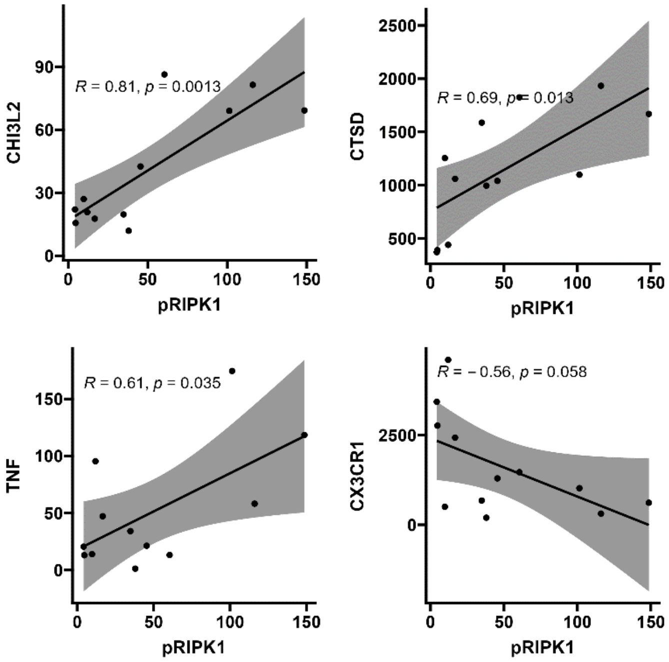


#### Astrocytes

C

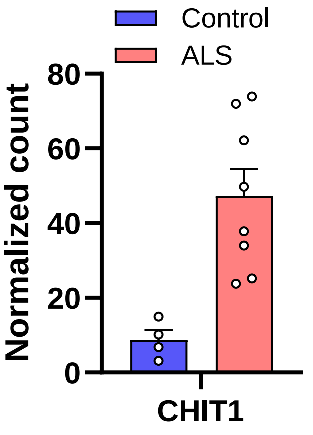


D

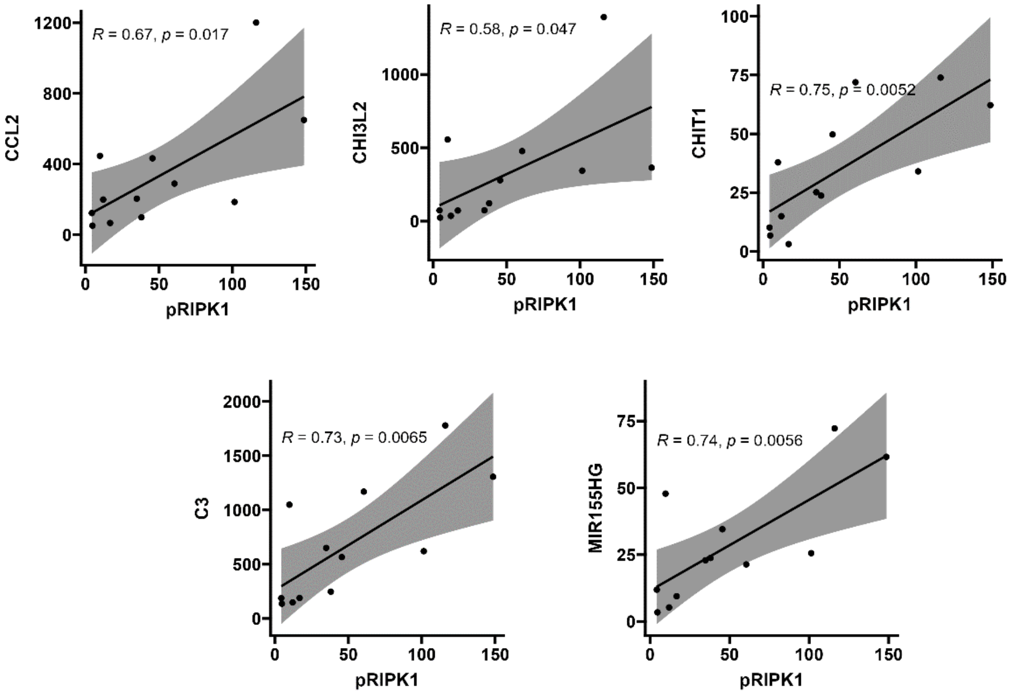

**Figure S6: Microglia and astrocyte DEGs correlate with RIPK1 kinase activation in human ALS spinal cords**

A-B) Pseudobulk DESeq2 normalized counts (A) and correlation graphs (B) depicting microglial gene expression correlating with pRIPK1 level in ALS (n=8) and control (n=4) spinal cords

C-D) Pseudobulk DESeq2 normalized counts (C) and correlation graphs (D) depicting astrocyte gene expression correlating with pRIPK1 level in ALS (n=8) and control (n=4) spinal cords

Data depict biological replicates and error bars represent mean ± SEM. Pearson correlation coefficient and two-tailed p-value were used (B, D).

### Figure S7:

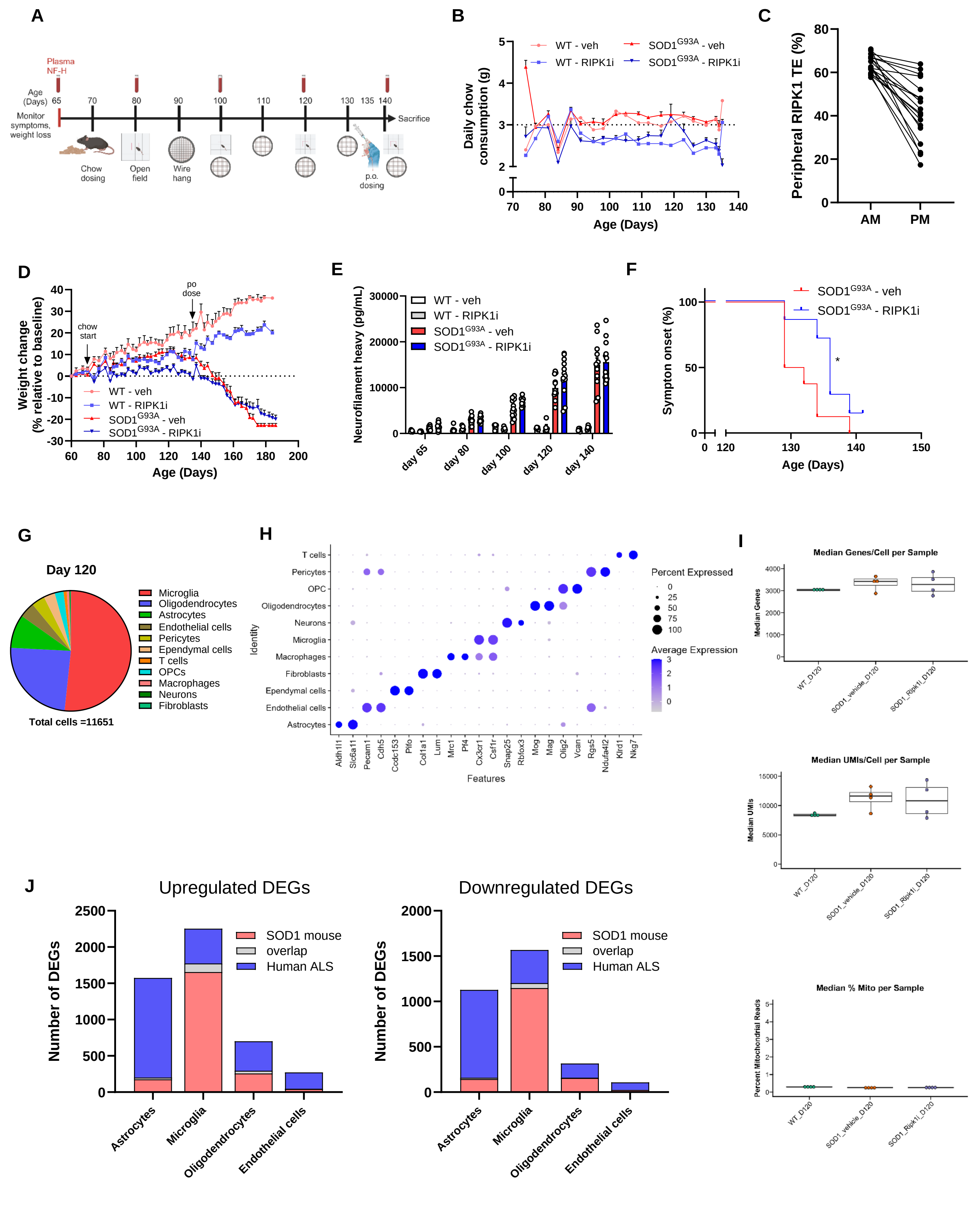
A
Plasma NF-H
Age (Days) 65 70 80 90 100 110 120 130 135 140
Monitor symptoms, weight loss
Sacrifice
Chow dosing
Open field
Wire hang
p.o. dosing
B
WT - veh
WT - RIPK1i
SOD1G93A - veh
SOD1G93A - RIPK1i
Daily chow consumption (g)
Age (Days)
C
Peripheral RIPK1 TE (%)
AM
PM
D
Weight change (% relative to baseline)
chow start
po dose
WT - veh
WT - RIPK1i
SOD1G93A - veh
SOD1G93A - RIPK1i
Age (Days)
E
Neurofilament heavy (pg/mL)
WT - veh
WT - RIPK1i
SOD1G93A - veh
SOD1G93A - RIPK1i
day 65
day 80
day 100
day 120
day 140
F
Sympton onset (%)
SOD1G93A - veh
SOD1G93A - RIPK1i
Age (Days)
G
Day 120
Microglia
Oligodendrocytes
Astrocytes
Endothelial cells
Pericytes
Ependymal cells
T cells
OPCs
Macrophages
Neurons
Fibroblasts
Total cells =11651
H
Identity
T cells
Pericytes
OPC
Oligodendrocytes
Neurons
Microglia
Macrophages
Fibroblasts
Ependymal cells
Endothelial cells
Astrocytes
Aldh1l1 Slc6a11 Pecam1 Cdh5 Ccdc153 Pifo Col1a1 Lum Mrc1 Pf4 Cx3cr1 Csf1r Snap25 Rbfox3 Mog Mag Olig2 Vcan Rgs5 Ndufa4l2 Klrd1 Nkg7
Features
Percent Expressed
Average Expression
I
Median Genes/Cell per Sample
Median Genes
Median UMIs/Cell per Sample
Median UMIs
Median % Mito per Sample
Percent Mitochondrial Reads
WT_D120
SOD1_vehicle_D120
SOD1_Ripk1i_D120
J
Upregulated DEGs
Downregulated DEGs
Number of DEGs
SOD1 mouse
overlap
Human ALS
Astrocytes
Microglia
Oligodendrocytes
Endothelial cells

**Figure S7: SOD1 mouse dosing and study scheme, single cell RNA-seq analysis of spinal cords**

A) Schematic demonstrating study design for RIPK1 inhibition in WT and $SOD1^{G93A}$ mice, including blood collection for plasma neurofilament and wire hang and open field activity tests, created with Biorender

B) Graph depicting daily amount of control or RIPK1 inhibitor chow consumed per WT or $SOD1^{G93A}$ mouse from postnatal day 70 to day 135 (n=5 WT control and WT RIPK1i mice in 1 cage each; n=12 $SOD1^{G93A}$ control and n=13 $SOD1^{G93A}$ RIPK1i mice in 4 cages each); estimated daily intake was 3g per mouse

C) Peripheral RIPK1 target engagement in the blood assessed by epitope masking assay in WT (n=5) and $SOD1^{G93A}$ (n=13) mice on postnatal day 121, after 50 days of chow dosing with RIPK1 inhibitor

D) Graph depicting the change from baseline weight for control (veh) or RIPK1 inhibitor (RIPK1i)-treated WT and $SOD1^{G93A}$ mice

E) Bar graph depicting plasma neurofilament heavy levels over time in vehicle or RIPK1 inhibitor-treated WT and $SOD1^{G93A}$ mice

F) Kaplan-Meier curve depicting symptom onset in a second cohort of $SOD1^{G93A}$ mice fed control (n=8) or RIPK1 inhibitor (n=7) chow. Mice (n=4) from the initial cohort of n=12/group were used for scRNA-seq

G) Pie chart quantifying cell types captured by scRNA-seq analysis of naïve WT, vehicle- and RIPK1 inhibitor-treated $SOD1^{G93A}$ mouse spinal cords isolated on postnatal day 120 (n=4 per group)

H) Dot plot showing feature genes for all major spinal cord cell types captured by scRNA-seq of naïve WT, vehicle- or RIPK1 inhibitor-treated $SOD1^{G93A}$ mice

I) Bar graphs depicting the median number of genes, unique molecular identifiers, and percent mitochondrial genes from scRNA-seq analysis of naïve WT, vehicle or RIPK1 inhibitor-treated $SOD1^{G93A}$ mouse spinal cords

J) Bar graphs depicting unique or overlapping upregulated or downregulated DEGs across various cell types in scRNA-seq analysis of vehicle-treated $SOD1^{G93A}$ mice and snRNA-seq analysis of human ALS cervical spinal cords relative to control

Data depict biological replicates and error bars represent mean ± SEM. Gehan-Breslow-Wilcoxon was performed (F). $*p < 0.05$

### Figure S8:

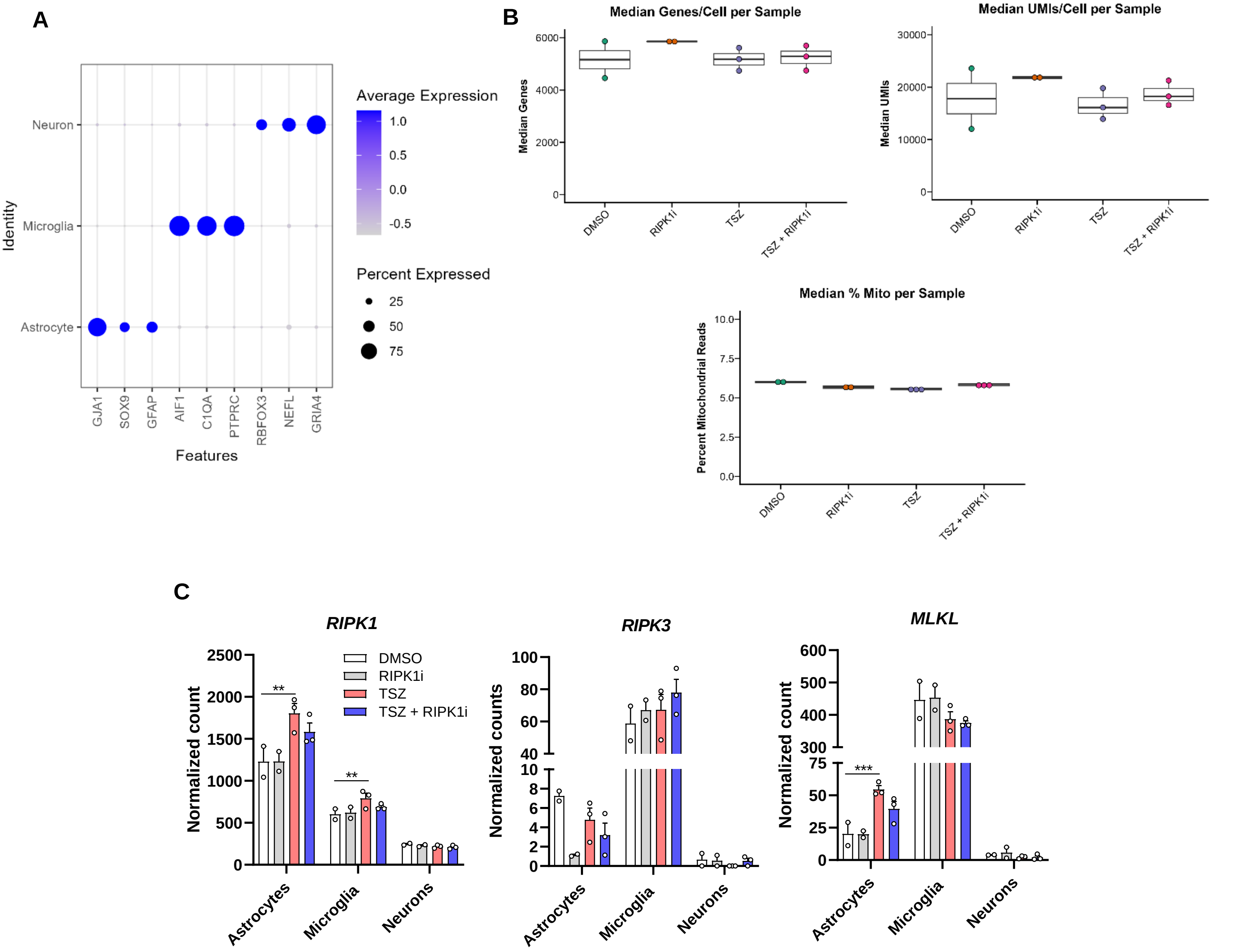


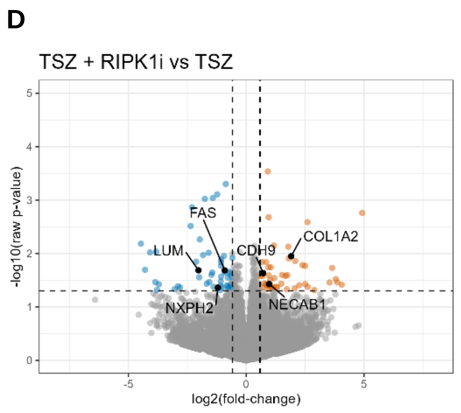


E

| Biological Process | pValue |
|---|---|
| regulation of immune system process | 2.14E-02 |
| forebrain development | 2.14E-02 |
| telencephalon development | 2.14E-02 |
| regulation of immune response | 4.98E-02 |
| leukocyte activation | 4.98E-02 |

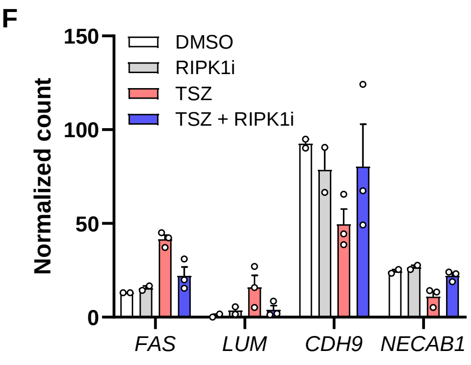

**Figure S8: Single nucleus RNA-seq analysis of iPSC tri-culture with neuronal gene expression analysis**

A) Dot plot depicting feature genes for astrocytes, microglia, and neurons captured via scRNA-seq in the TSZ-treated iPSC tri-culture

B) Bar graphs depicting median number of genes, unique molecular identifiers, and percent mitochondrial genes from scRNA-seq analysis of iPSC tri-culture treated with TSZ in the presence or absence of a RIPK1 inhibitor (n=2 DMSO, RIPK1i; n=3 TSZ, TSZ + RIPK1i)

C) Pseudobulk DESeq2 normalized counts for *RIPK1*, *RIPK3*, and *MLKL* expression in astrocytes, microglia, or neurons from the TSZ-stimulated tri-culture

D) Volcano plot showing neuronal DEGs in tri-culture samples stimulated with TSZ versus TSZ and RIPK1 inhibitor (TSZ + RIPK1i); (cutoff $|FC| > 1.5$, $p < 0.05$)

E) Gene Ontology terms for biological processes for RIPK1-dependent neuronal DEGs in the TSZ-stimulated tri-culture

F) Pseudobulk DESeq2 normalized counts highlighting examples of neuronal RIPK1-dependent gene expression in the TSZ-stimulated tri-culture

Data depict technical replicates from a tri-culture and error bars represent mean ± SEM. FDR (Benjamini-Hochberg) was used (E). **$p < 0.01$, ***$p < 0.001$; T, TNF; S, Smac mimetic; Z, zVAD-fmk
